# Rapalogs downmodulate intrinsic immunity and promote cell entry of SARS-CoV-2

**DOI:** 10.1101/2021.04.15.440067

**Authors:** Guoli Shi, Abhilash I. Chiramel, Tiansheng Li, Kin Kui Lai, Adam D. Kenney, Ashley Zani, Adrian Eddy, Saliha Majdoul, Lizhi Zhang, Tirhas Dempsey, Paul A. Beare, Swagata Kar, Jonathan W. Yewdell, Sonja M. Best, Jacob S. Yount, Alex A. Compton

## Abstract

SARS-CoV-2 infection in immunocompromised individuals is associated with prolonged virus shedding and evolution of viral variants. Rapamycin and its analogs (rapalogs, including everolimus, temsirolimus, and ridaforolimus) are FDA-approved as mTOR inhibitors for the treatment of human diseases, including cancer and autoimmunity. Rapalog use is commonly associated with increased susceptibility to infection, which has been traditionally explained by impaired adaptive immunity. Here, we show that exposure to rapalogs increases susceptibility to SARS-CoV-2 infection in tissue culture and in immunologically naive rodents by antagonizing the cell-intrinsic immune response. By identifying one rapalog (ridaforolimus) that is less potent in this regard, we demonstrate that rapalogs promote Spike-mediated entry into cells by triggering the degradation of antiviral proteins IFITM2 and IFITM3 via an endolysosomal remodeling program called microautophagy. Rapalogs that increase virus entry inhibit the mTOR-mediated phosphorylation of the transcription factor TFEB, which facilitates its nuclear translocation and triggers microautophagy. In rodent models of infection, injection of rapamycin prior to and after virus exposure resulted in elevated SARS-CoV-2 replication and exacerbated viral disease, while ridaforolimus had milder effects. Overall, our findings indicate that preexisting use of certain rapalogs may elevate host susceptibility to SARS-CoV-2 infection and disease by activating lysosome-mediated suppression of intrinsic immunity.

**Significance:** Rapamycin is an immunosuppressant used in humans to treat cancer, autoimmunity, and other disease states. Here, we show that rapamycin and related compounds promote the first step of the SARS-CoV-2 infection cycle—entry into cells—by disarming cell-intrinsic immune defenses. We outline the molecular basis for this effect by identifying a rapamycin derivative that is inactive, laying the foundation for improved mTOR inhibitors that do not suppress intrinsic immunity. We find that rapamycin analogs that promote SARS-CoV-2 entry are those that activate TFEB, a transcription factor that triggers the degradation of antiviral membrane proteins inside of cells. Finally, rapamycin administration to rodents prior to SARS-CoV-2 challenge results in enhanced viral disease, revealing that its use in humans may increase susceptibility to infection.

## Introduction

Severe acute respiratory syndrome coronavirus 2 (SARS-CoV-2) emerged in humans in 2019 following a species jump from bats and is the cause of COVID-19, a respiratory and multi-organ disease of variable severity (1, 2). The characterization of virus-host interactions that dictate SARS-CoV-2 infection and COVID-19 severity is a major priority for public health (3). Immune impairment, such as that resulting from cancer, has been associated with prolonged SARS-CoV-2 shedding, the seeding of “super-spreader” events, and the evolution of viral variants (4–8).

One group of compounds being considered for the treatment of COVID-19-related immunopathology are rapamycin (sirolimus, Rapamune) and rapamycin analogs (rapalogs) (9–20). As Food and Drug Administration-approved inhibitors of mammalian target of rapamycin (mTOR) kinase, these macrolide compounds are used therapeutically to inhibit the processes of cancer, autoimmunity, graft versus host disease, atherosclerosis, and aging (21). Rapalogs, including everolimus (RAD-001), temsirolimus (Torisel, CCI-779), and ridaforolimus (deforolimus, AP-23573), were developed to decrease the half-life of rapamycin in vivo in order to minimize the systemic immunosuppression caused by rapamycin use, which is associated with increased susceptibility to infections (22–26). Differing by only a single functional group at carbon-40 **(Figure 1)**, it is believed that rapamycin and rapalogs share the same molecular mechanism of action to inhibit mTOR kinase—they bind to FK506-binding proteins (FKBP) and the resulting complex physically interacts with mTOR and disrupts its signaling (25, 27).

**Figure 1:**
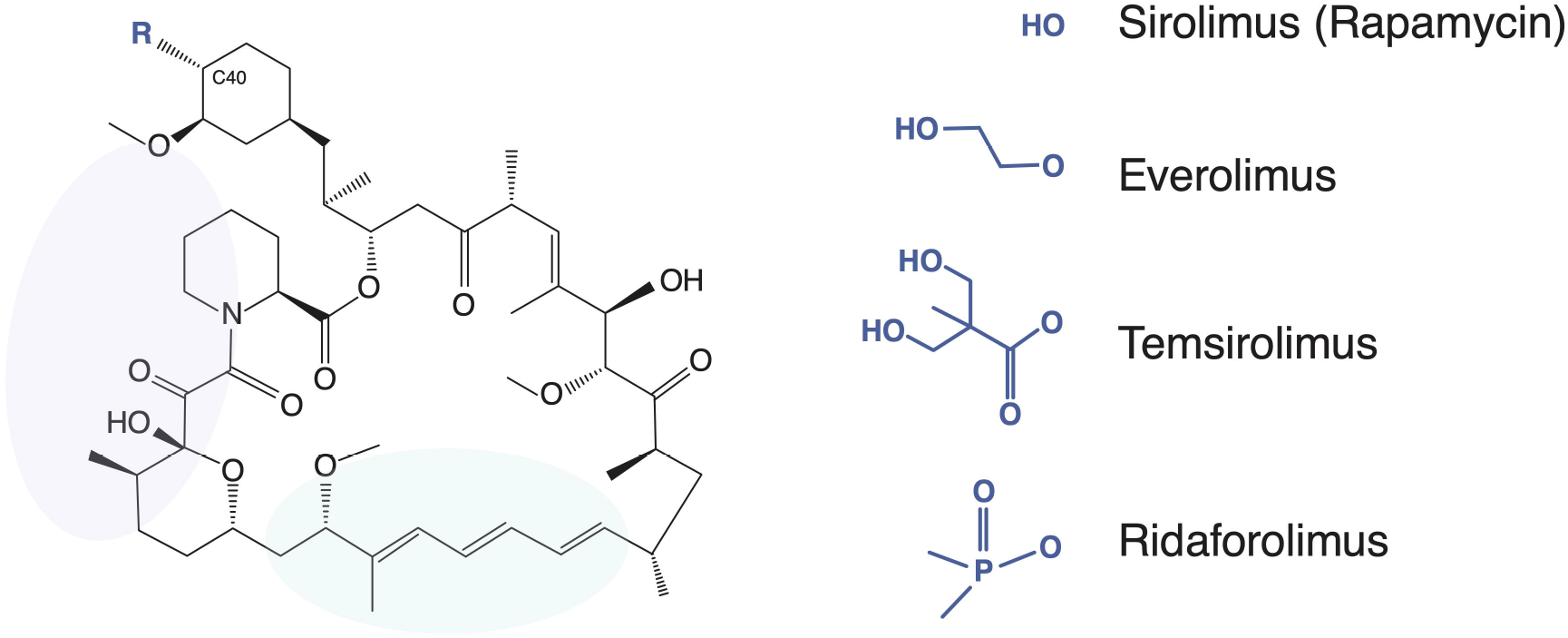
Rapamycin and its analogs share a macrolide structure but differ by the functional group present at carbon-40. Violet and green bubbles indicate the FKBP- and mTOR-binding sites, respectively.

Activation of mTOR promotes cell growth, cell proliferation, and cell survival (28). In addition, mTOR activation promotes pro-inflammatory T-cell differentiation and mTOR inhibitors have been used to block lymphocyte proliferation and cytokine storm (29). Since respiratory virus infections like SARS-CoV-2 can cause disease by provoking hyper-inflammatory immune responses that result in immunopathology (30–32), rapalogs are being tested as treatments to decrease viral disease burden. At least three active clinical trials have been designed to test the impact of rapamycin on COVID-19 severity in infected patients (NCT04461340, NCT04341675, NCT04371640).

In addition to their potential utility for mitigating disease in individuals already infected by SARS-CoV-2, there are also calls to use rapalogs as antiviral agents to inhibit virus infection itself (i.e. as a prophylactic) (33). It was recently shown that rapalogs inhibit SARS-CoV-2 replication when added to cells post-infection (34), attesting to a potential use of rapalogs as antivirals in infected individuals. Nonetheless, rapalogs are known to induce an immunosuppressed state in humans characterized by an increased rate of infections, including those caused by respiratory viruses. Furthermore, rapamycin administration concurrent with virus challenge has been shown to promote Influenza A replication in mice and to exacerbate viral disease (35, 36), but the mechanism was unknown. We previously found that exposure of human and murine cells to rapamycin induced the lysosomal degradation of a select group of cellular proteins, including the interferon-inducible transmembrane (IFITM) proteins, and rendered cells more permissive to infection by Influenza A virus and gene-delivering lentiviral vectors (37, 38). IFITM1, IFITM2, and IFITM3 are expressed constitutively in a variety of tissues, are further upregulated by type-I, type-II, and type-III interferons, and are important components of cell-intrinsic immunity, the antiviral network that defends individual cells against virus invasion (39, 40). Nonetheless, it remained to be determined how rapamycin-mediated regulation of intrinsic immunity impacts host susceptibility to virus infection in vivo.

In this report, we show that rapalogs differentially counteract the constitutive and interferon-induced antiviral state in lung cells and increase permissiveness to SARS-CoV-2 infection. We found that the enhancing effect of rapalogs on SARS-CoV-2 infection is functionally linked to their capacity to trigger degradation of IFITM proteins, particularly IFITM2 and IFITM3. By identifying a rapalog that lacks this activity, we found that IFITM protein turnover and SARS-CoV-2 infection enhancement are associated with activation of TFEB, a master regulator of lysosome function that is regulated by mTOR. Administration of rapamycin to naive rodents four hours prior to experimental SARS-CoV-2 infection increased virus replication and viral disease severity, indicating for the first time that suppression of intrinsic immunity by rapamycin contributes to its immunosuppressive properties in vivo.

## Results

### Select rapalogs promote SARS-CoV-2 infection and downmodulate IFITM proteins in lung cells

To assess how rapamycin and rapalogs impact SARS-CoV-2 infection, we took advantage of a pseudovirus system based on human immunodeficiency virus (HIV). This pseudovirus (HIV-CoV-2) is limited to a single round of infection, cell entry is mediated by SARS-CoV-2 Spike, and infection of target cells is measured by luciferase activity. SARS-CoV-2 can enter cells via multiple routes, and sequential proteolytic processing of Spike is essential to this process. SARS-CoV-2 Spike is cleaved at a polybasic motif (RRAR) located at the S1/S2 boundary by furin-like proteases in virus-producing cells prior to release. Subsequently, the S2’ site is cleaved by the trypsin-like protease TMPRSS2 on the target cell surface or by cathepsins B and L in target cell endosomes, triggering membrane fusion at those sites (41–43).

We previously found that a four-hour pre-treatment of cells with 20 μM quantities of rapamycin triggered the degradation of human IFITM3 and enhanced cellular susceptibility to Influenza A virus infection (38). Therefore, we pre-treated A549-ACE2 (transformed human lung epithelial cells that overexpress the SARS-CoV-2 receptor, human ACE2) with 20 μM rapamycin, everolimus, temsirolimus, ridaforolimus, or DMSO (vehicle control) for four hours and then challenged cells with HIV-CoV-2. Interestingly, we found that rapalogs promoted Spike-mediated infection to different extents: rapamycin, everolimus, and temsirolimus significantly enhanced infection (up to 5-fold) while ridaforolimus did not **(Figure 2A)**. To determine whether rapalogs promote cell permissiveness to infection by upregulating dependency factors or by downregulating restriction factors, we performed the same experiment in cells pre-treated with type-I interferon. While type-I interferon suppressed infection by approximately 90%, the addition of rapamycin, everolimus, and temsirolimus resulted in rescue of infection by up to 20-fold **(Figure 2A)**. As a result, infection levels were partially restored to those achieved in the absence of interferon, with everolimus having the greatest boosting effect and ridaforolimus, the least. These results indicate that rapalogs differentially promote SARS-CoV-2 Spike-mediated infection by counteracting intrinsic antiviral defenses in lung cells to different extents.

**Figure 2:**
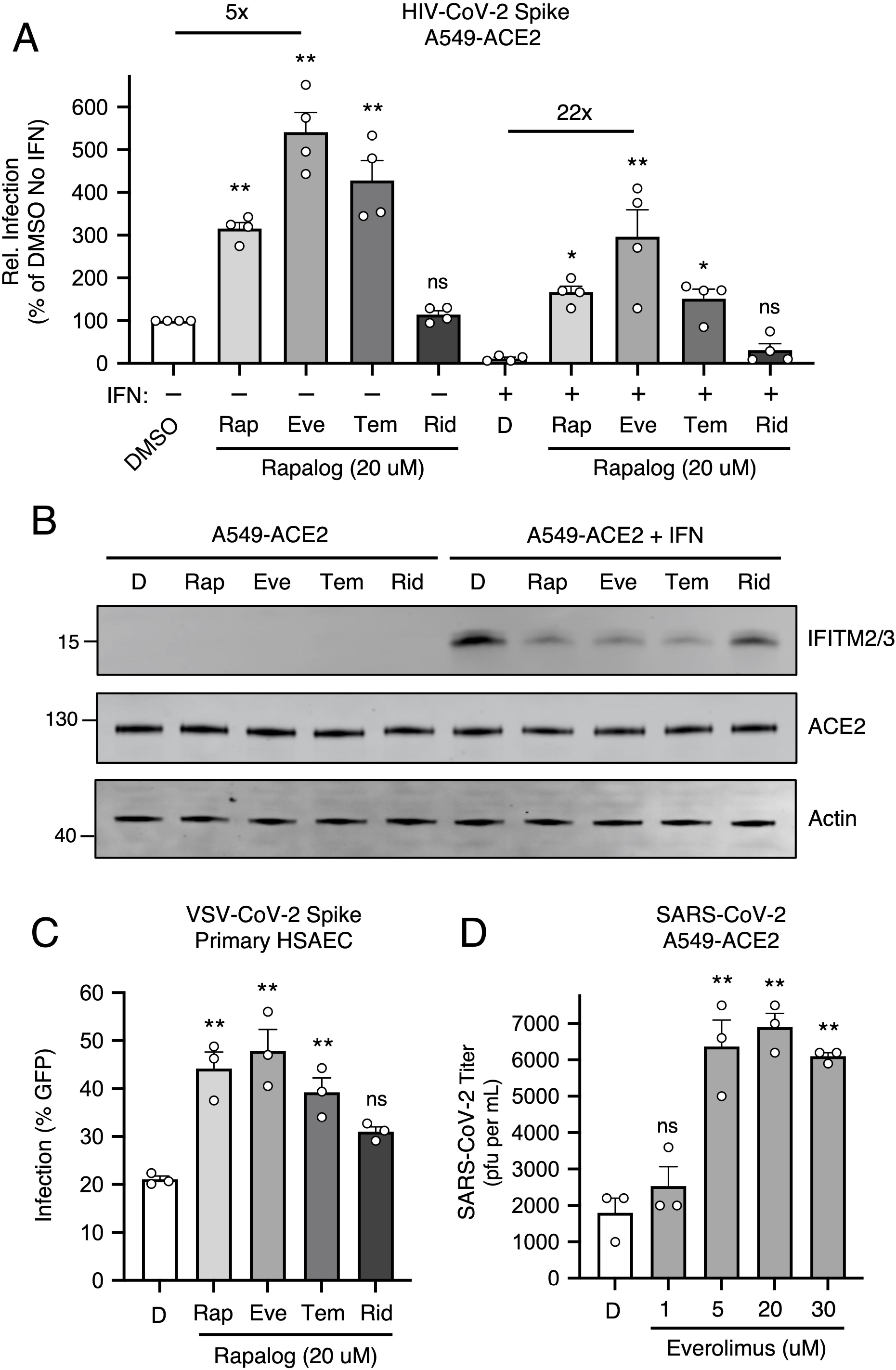
Rapalogs promote SARS-CoV-2 infection in lung epithelial cells to different extents by counteracting the intrinsic antiviral state. (A) A549-ACE2 were treated with or without type I interferon (250 U/mL) for 18 hours and then treated with 20 µM rapamycin (Rap), everolimus (Eve), temsirolimus (Tem), ridaforolimus (Rid), or an equivalent volume of DMSO (D) for 4 hours. HIV-CoV-2 (100 ng p24 equivalent) was added to cells and infection was measured by luciferase activity at 48 hours post-infection. Luciferase units were normalized to 100 in the DMSO condition in the absence of interferon. (B) A549-ACE2 cells from (A) were subjected to SDS-PAGE and Western blot analysis. Immunoblotting was performed with anti-IFITM2/3, anti-ACE2, and anti-actin (in that order) on the same nitrocellulose membrane. Numbers and tick marks indicate size (kilodaltons) and position of protein standards in ladder. (C) Primary HSAEC were treated with 20 µM Rap, Eve, Tem, Rid, or an equivalent volume of DMSO for 4 hours. VSV-CoV-2 (50 µL) was added to cells and infection was measured by GFP expression at 24 hours post-infection using flow cytometry. (D) A549-ACE2 were treated with varying concentrations of Eve or DMSO (equivalent to 30 µM of Eve) for 4 hours. SARS-CoV-2 (nCoV-WA1-2020; MN985325.1) was added to cells at an MOI of 0.1 and infectious titers were measured in VeroE6 cells by calculating the TCID_50_ per mL of supernatants recovered at 24 hours post-infection. TCID_50_ (pfu per mL) values are shown. Means and standard error were calculated from 3-4 experiments. Statistical analysis was performed with one-way ANOVA and asterisks indicate significant difference from DMSO. *, p < 0.05; **, p < 0.01. Rel.; relative. pfu; plaque forming units.

Type-I interferon treatment of A549-ACE2 cells resulted in upregulation of IFITM2 and IFITM3, as detected by an antibody recognizing both proteins in whole cell lysates **(Figure 2B)**. A549-ACE2 cells express low but detectable levels of IFITM2/3 in the absence of interferon treatment **(Supplemental Figure 1A)**. Consistent with our previous publication, addition of rapamycin resulted in substantial loss of IFITM2/3 protein levels from cells. In a manner that mirrored the differential effects of rapalogs on pseudovirus infection, everolimus and temsirolimus greatly diminished IFITM2/3 levels while ridaforolimus reduced IFITM2/3 to a lesser extent **(Figure 2B and Supplemental Figure 1A)**. In contrast, ACE2 levels were not affected by interferon nor by rapalog treatment. Therefore, rapamycin derivatives may facilitate infection by antagonizing constituents of intrinsic immunity, including IFITM2/3, and this activity is determined by the chemical moiety found at carbon-40 of the macrolide structure.

To extend our findings to primary lung cells, we performed similar experiments in human small airway epithelial cells (HSAEC). While these cells were not permissive to HIV-CoV-2, they were susceptible to infection by pseudovirus based on vesicular stomatitis virus (VSV-CoV-2) whereby infection is reported by GFP expression. Pre-treatment of HSAEC with rapalogs enhanced VSV-CoV-2 infection to varying extents, but as observed in A549-ACE2 cells, everolimus exhibited the greatest effect and ridaforolimus, the least. Endogenous IFITM3 was readily detected in HSAEC under basal conditions (in the absence of interferon) and its levels were downmodulated differentially by rapalogs. However, IFITM1 was barely detected and IFITM2 was not detected at all. **(Supplemental Figure 1B)**. siRNA-mediated knockdown of IFITM3 in HSAEC resulted in enhanced VSV-CoV-2 infection, indicating that IFITM3 restricts Spike-mediated infection in these cells **(Supplemental Figure 1C)**. We also treated transformed nasal epithelial cells (UNCNN2TS) with rapalogs in order to assess an impact on endogenous IFITM3 levels. As observed in HSAEC, downmodulation of IFITM3 occurred following treatment of UNCNN2TS with rapamycin, everolimus, temsirolimus, and to a lesser extent, ridaforolimus **(Supplemental Figure 1D)**.

Since 20 μM quantities of rapalogs promoted pseudovirus infection mediated by SARS-CoV-2 Spike, we tested how pretreatment of A549-ACE2 cells with varying amounts of everolimus impacted infection by replication-competent SARS-CoV-2. We observed a dose-dependent enhancement of infectious SARS-CoV-2 yield in supernatants of infected cells (up to 4-fold) **(Figure 2D)**. Therefore, everolimus boosts pseudovirus infection and SARS-CoV-2 infection to similar extents, and since Spike is the only viral component shared between the two sources of infection, cellular entry is the infection stage inhibited by the intrinsic defenses that are sensitive to downmodulation by rapalogs.

### Rapalogs facilitate cell entry mediated by various viral fusion proteins

In order to gain a greater mechanistic understanding of the effects of rapalogs on SARS-CoV-2 infection, we took advantage of HeLa cells overexpressing ACE2 (HeLa-ACE2). HeLa-ACE2 were pre-treated for four hours with increasing amounts of everolimus and then challenged with SARS-CoV-2. Everolimus increased titers of infectious virus released into supernatants in a dose-dependent manner, and to a greater extent than was observed for A549-ACE2 cells **(Figure 3A)**. Furthermore, we found that pre-treatment of cells with 20 μM amounts of rapalogs enhanced SARS-CoV-2 titers to varying extents—rapamycin, everolimus, and temsirolimus significantly boosted SARS-CoV-2 infection (up to 10-fold), while ridaforolimus had less of an impact **(Figure 3B)**. We also performed infections of HeLa-ACE2 with HIV-CoV-2 pseudovirus, and the results were similar: the impact of ridaforolimus was minimal while the other three compounds significantly boosted Spike-mediated infection **(Figure 3C)**. To test the link between infection enhancement and downmodulation of IFITM proteins by rapalogs, we probed for levels of IFITM3, IFITM2, and IFITM1 by immunoblotting whole cell lysates using specific antibodies. All IFITM proteins were readily detected in HeLa-ACE2 in the absence of interferon. IFITM3, IFITM2, and IFITM1 were significantly downmodulated following treatment with rapamycin, everolimus, and temsirolimus **(Figure 3D)**. Levels of IFITM3 were quantified over multiple experiments and presented as an average. The results show that all rapalogs led to significant decreases in IFITM3 protein, but ridaforolimus was least potent in this regard **(Figure 3E)**. The loss of IFITM2/3 protein was confirmed by confocal immunofluorescence microscopy of intact cells (**Figure 3F**). Furthermore, prolonged treatment (24 hours) of cells with everolimus and temsirolimus resulted in prolonged suppression of IFITM2 and IFITM3 protein levels **(Supplemental Figure 2A)**. In contrast, ACE2 levels and ACE2 subcellular distribution were unaffected by rapalog treatment **(Figure 3D and Supplemental Figure 2B)**. Furthermore, rapalogs did not significantly decrease cell viability under the conditions tested **(Supplemental Figure 2C)**.

**Figure 3:**
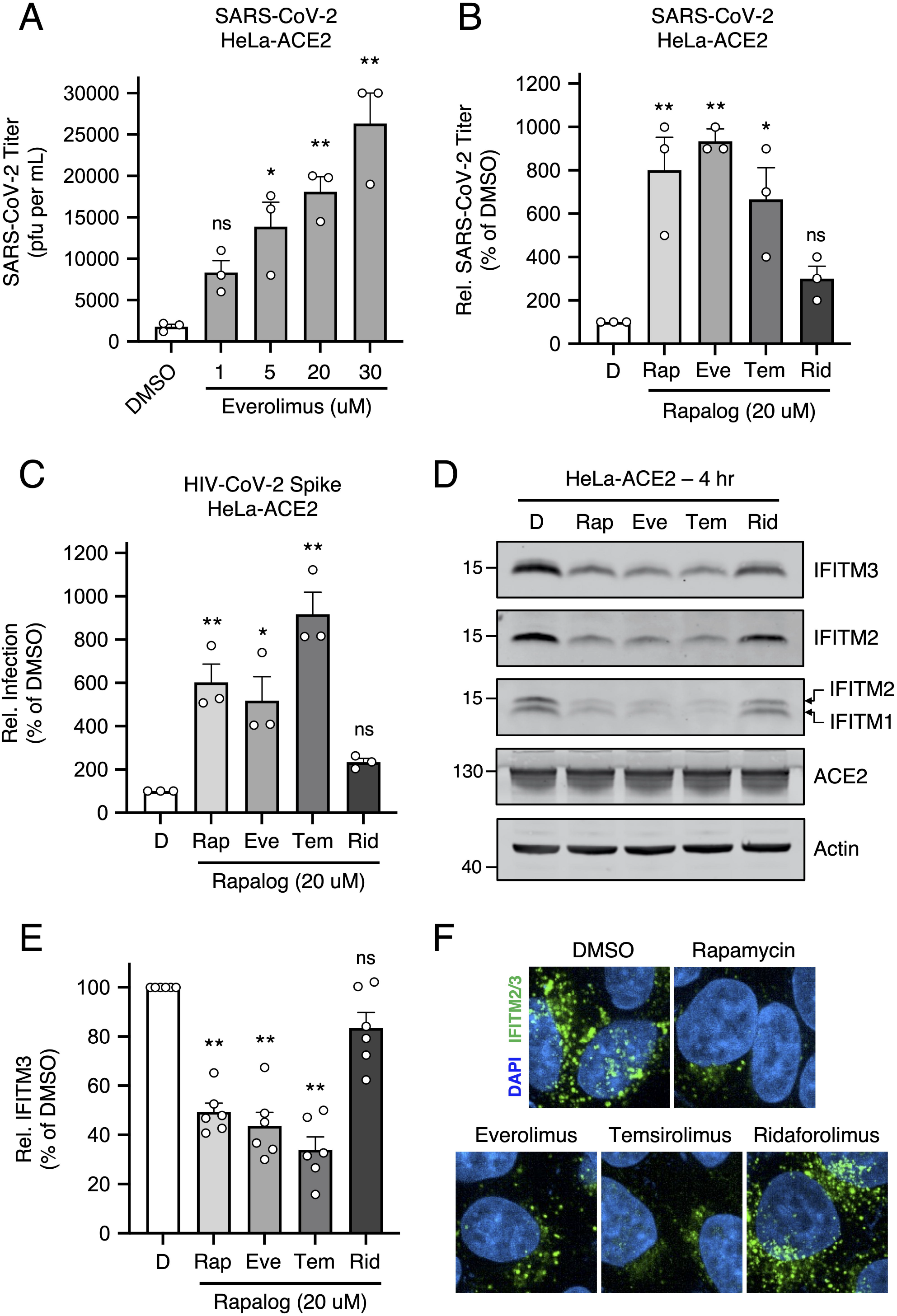
Rapalogs promote SARS-CoV-2 infection in HeLa-ACE2 cells. (A) HeLa-ACE2 were treated with varying concentrations of Eve or DMSO for 4 hours. SARS-CoV-2 (nCoV-WA1-2020; MN985325.1) was added to cells at MOI 0.1 and infectious titers were measured in VeroE6 cells by calculating the TCID_50_ of supernatants recovered at 24 hours post-infection. TCID_50_ (pfu per mL) values are shown. (B) HeLa-ACE2 were treated with 20 µM Rap, Eve, Tem, Rid, or an equivalent volume of DMSO for 4 hours. SARS-CoV-2 (nCoV-WA1-2020; MN985325.1) was added to cells at MOI 0.1 and infectious titers were measured in VeroE6 cells by calculating the TCID_50_ per mL of supernatants recovered at 24 hours post-infection. TCID_50_ per mL values were normalized to 100 in the DMSO condition. (C) HeLa-ACE2 were treated with 20 µM Rap, Eve, Tem, Rid, or an equivalent volume of DMSO for 4 hours. HIV-CoV-2 (100 ng p24 equivalent) was added to cells and infection was measured by luciferase activity at 48 hours post-infection. Luciferase units were normalized to 100 in the DMSO condition. (D) HeLa-ACE2 cells from (C) were subjected to SDS-PAGE and Western blot analysis. Immunoblotting was performed with anti-IFITM2, anti-IFITM1, anti-IFITM3, anti-ACE2, and anti-actin (in that order) on the same nitrocellulose membrane. (E) IFITM3 levels from (D) were normalized to actin levels and summarized from 5 independent experiments. (F) HeLa-ACE2 were treated with 20 µM Rap, Eve, Tem, Rid, or an equivalent volume of DMSO for 4 hours and cells were fixed, stained with DAPI and anti-IFITM2/3, and imaged by confocal immunofluorescence microscopy. Images represent stacks of 5 Z-slices and one representative image is shown per condition. Means and standard error were calculated from 3-6 experiments. Statistical analysis was performed with one-way ANOVA and asterisks indicate significant difference from DMSO. *, p < 0.05; **, p < 0.01. Rel.; relative. pfu; plaque forming units.

We previously showed that lysosomal degradation of IFITM3 triggered by rapamycin requires endosomal complexes required for transport (ESCRT) machinery and multivesicular body (MVB)-lysosome fusion (38). We confirmed that depletion of IFITM proteins by rapalogs occurs at the post-translational level and requires endolysosomal acidification, since bafilomycin A1 prevented their loss **(Supplemental Figure 3A-B)**. The process by which rapalogs trigger IFITM protein degradation resembles endolysosomal microautophagy, an autophagy pathway that does not require an autophagosome intermediate (44–46). Treatment of cells with U18666A, an inhibitor of MVB formation and microautophagy, mostly prevented IFITM3 turnover in the presence of rapalogs **(Supplementary Figure 3B)**. In contrast, a selective inhibitor of vps34/PI3KC3 (essential for macroautophagy induction) did not **(Supplemental Figure 3C-D)**. Therefore, rapamycin and specific rapalogs trigger the degradation of endogenous factors mediating intrinsic resistance to SARS-CoV-2 infection, including the IFITM proteins, by promoting their turnover in lysosomes via endolysosomal microautophagy.

Enveloped virus entry into cells is a concerted process involving virus attachment to the cell surface followed by fusion of cellular and viral membranes. Since IFITM proteins are known to inhibit virus-cell membrane fusion, we quantified the terminal stage of HIV-CoV-2 entry by tracking the cytosolic delivery of beta-lactamase (BlaM) in single cells. We found that treatment of cells with rapamycin, everolimus, and temsirolimus resulted in enhanced HIV-CoV-2 entry while ridaforolimus was less impactful **(Figure 4A)**. To measure whether rapalogs promote the cell entry process driven by other coronavirus Spike proteins, we produced HIV incorporating Spike from SARS-CoV (HIV-CoV-1) or MERS-CoV (HIV-MERS-CoV). Infections by both HIV-CoV-1 and HIV-MERS-CoV were elevated by rapalog treatment in HeLa-ACE2 and HeLa-DPP4 cells, respectively, although the extent of enhancement was lower than that observed with HIV-CoV-2 **(Figure 4B-C)**. Consistently, ridaforolimus was the least active among the rapalogs tested and it did not significantly promote pseudovirus infection. Since we previously showed that rapamycin enhanced the cellular entry of Influenza A virus and VSV-G pseudotyped lentiviral vectors (38), we also assessed infection of pseudoviruses incorporating hemagglutinin (HIV-HA) or VSV G (HIV-VSV G). Rapamycin, everolimus, and especially temsirolimus boosted HA- and VSV G-mediated infections (up to 30-fold and 11-fold, respectively) **(Figure 4D-E)**. Since IFITM proteins have been previously shown to inhibit infection by SARS-CoV, MERS-CoV, VSV, and Influenza A virus (40), these data suggest that rapalogs promote infection, at least in part, by lowering the barrier to virus entry imposed by IFITM proteins.

**Figure 4:**
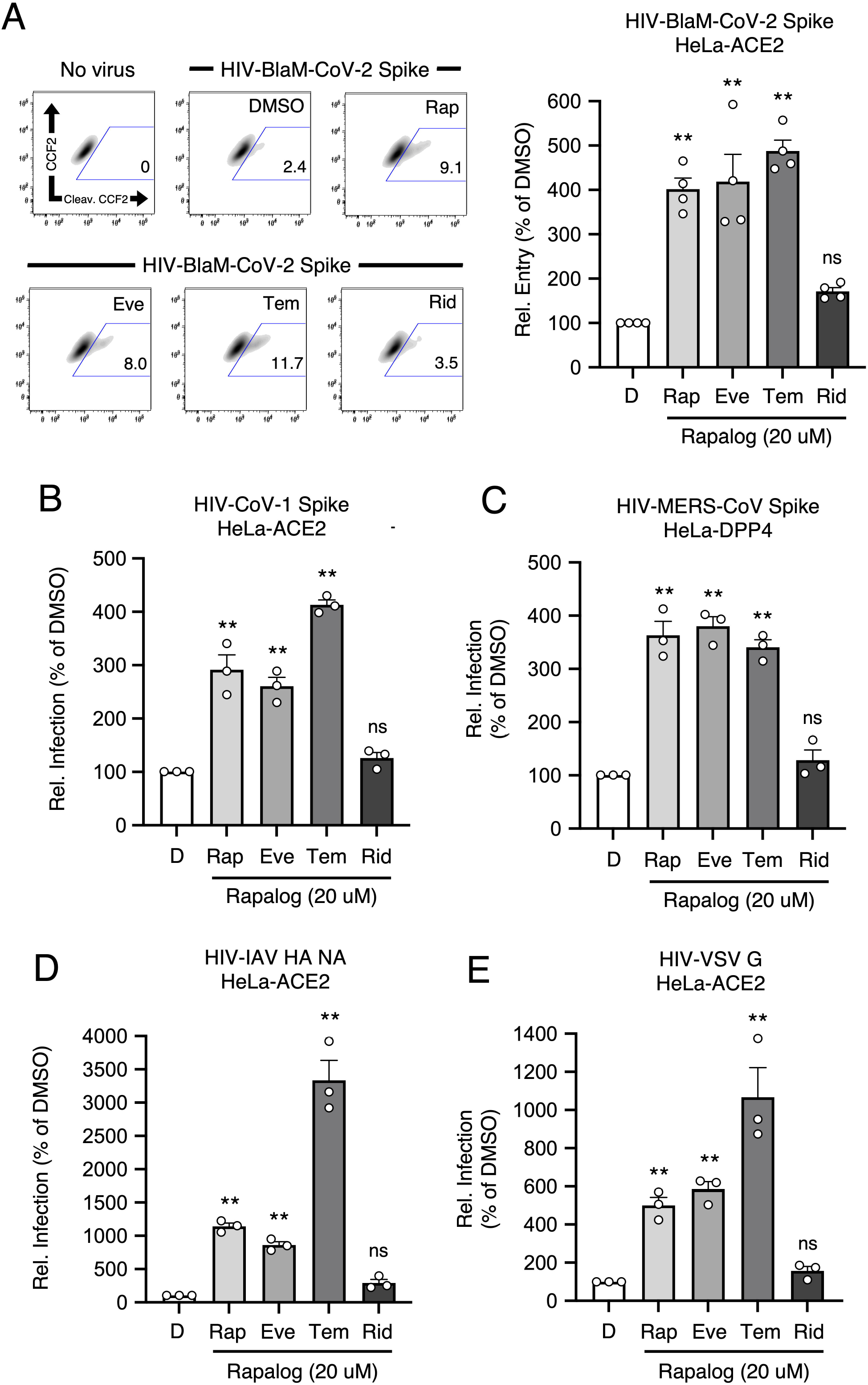
Rapalogs promote cell entry mediated by diverse viral fusion proteins. (A) HeLa-ACE2 were treated with 20 µM Rap, Eve, Tem, Rid, or an equivalent volume of DMSO for 4 hours. HIV-CoV-2 S pseudovirus incorporating BlaM-Vpr (HIV-BlaM-CoV-2) was added to cells for 2 hours and washed. Cells were incubated with CCF2-AM for an additional 2 hours and fixed. Cleaved CCF2 was measured by flow cytometry. Dot plots visualized as density plots from one representative experiment are shown on the left and the percentage of CCF2+ cells which exhibit CCF2 cleavage is indicated. Summary data representing the average of four experiments is shown on the right. (B) HIV-CoV-1, (C) HIV-MERS-CoV, (D) HIV-IAV HA, or (E) HIV-VSV G were added to HeLa-ACE2 or HeLa-DPP4 cells as in (A) and infection was measured by luciferase activity at 48 hours post-infection. Luciferase units were normalized to 100 in the DMSO condition. Means and standard error were calculated from 3-4 experiments. Statistical analysis was performed with one-way ANOVA and asterisks indicate significant difference from DMSO. *, p < 0.05; **, p < 0.01. Rel.; relative.

### IFITM2/3 mediate the rapalog-sensitive barrier to SARS-CoV-2 infection in HeLa-ACE2

To formally test the link between rapalog-mediated depletion of IFITM proteins and entry by SARS-CoV-2 Spike, we used HeLa cells in which IFITM1, IFITM2, and IFITM3 were knocked out (*IFITM1-3* KO) and introduced human ACE2 by transient transfection **(Figure 5A)**. IFITM2 alone or IFITM2 and IFITM3 were restored in *IFITM1-3* KO cells by transient overexpression **(Figure 5B)** and cells were challenged with HIV-CoV-2. Relative to WT cells, HIV-CoV-2 infection was approximately 50-fold higher in *IFITM1-3* KO cells, indicating that endogenous IFITM proteins restrict SARS-CoV-2 Spike-mediated infection in this cell type. Furthermore, while temsirolimus significantly promoted infection by 10-fold in WT cells, little to no enhancement was observed in *IFITM1-3* KO cells **(Figure 5C)**. Ectopic expression of IFITM2 inhibited infection and partially restored sensitivity to temsirolimus, while the combination of IFITM2 and IFITM3 restricted infection further and fully restored temsirolimus sensitivity. These findings indicate that temsirolimus promotes Spike-mediated infection in HeLa-ACE2 cells by lowering levels of endogenous IFITM2 and IFITM3. In accordance with the role played by endosomal IFITM2/3 in protecting cells against SARS-CoV-2 infection (47), pseudovirus infection mediated by Omicron (BA.1) Spike (which favors the endosomal route for entry ((48)) was as sensitive to temsirolimus-mediated enhancement as infection mediated by ancestral (WA1) Spike (**Figure 5D**). These results suggest that select rapalogs promote SARS-CoV-2 infection by negating the antiviral action of IFITM2/3 in endosomes.

**Figure 5:**
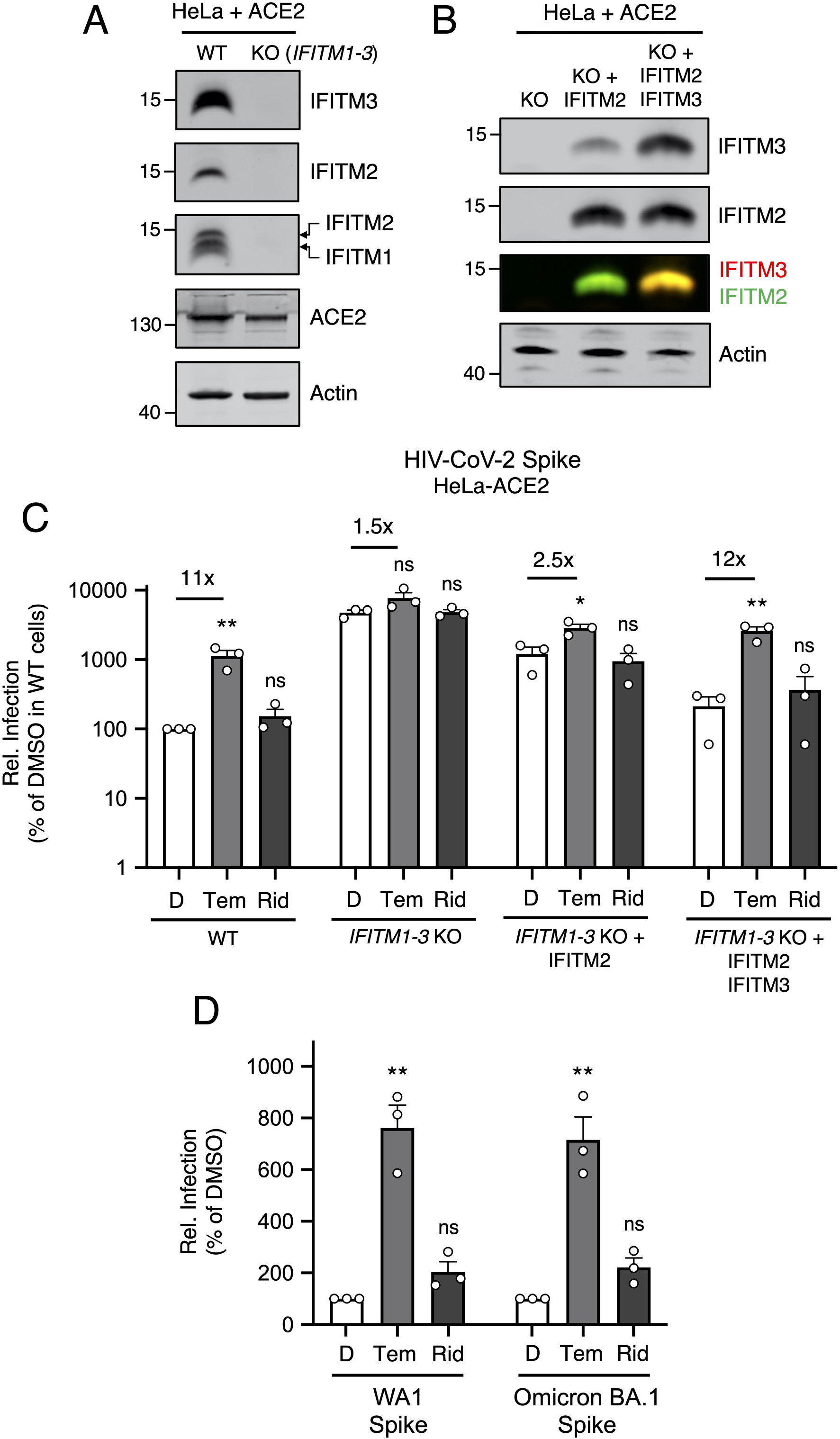
Select rapalogs enhance Spike-mediated infection in HeLa-ACE2 by inhibiting IFITM2 and IFITM3. (A) HeLa WT and HeLa *IFITM1-3* KO cells were transiently transfected with 0.150 µg pcDNA3.1-hACE2 for 24 hours. Whole cell lysates were subjected to SDS-PAGE and Western blot analysis. Immunoblotting was performed with anti-IFITM2, anti-IFITM3, anti-IFITM1, anti-ACE2, and anti-actin (in that order) on the same nitrocellulose membrane. (B) HeLa *IFITM1-3* KO were transfected with IFITM2 or IFITM2 and IFITM3 and SDS-PAGE and Western blot analysis was performed. (C) HIV-CoV-2 was added to transfected cells from (B) and infection was measured by luciferase activity at 48 hours post-infection. Luciferase units were normalized to 100 in HeLa WT cells treated with DMSO. (D) HeLa WT were transiently transfected with 0.150 µg pcDNA3.1-hACE2 for 24 hours. HIV-CoV-2 decorated with ancestral Spike (WA1) or Omicron Spike (BA.1) was added and infection was measured by luciferase activity at 48 hours post-infection. Luciferase units were normalized to 100 in cells treated with DMSO for both pseudoviruses. Means and standard error were calculated from 3 experiments. Statistical analysis was performed with one-way ANOVA and asterisks indicate significant difference from nearest DMSO condition. *, p < 0.05; **, p < 0.01. ns; not significant. Rel.; relative.

Since human IFITM proteins have been reported to promote SARS-CoV-2 infection in certain cell types, including the lung epithelial cell line Calu-3 (49), we tested the impact of rapalogs on HIV-CoV-2 infection in this cell type. Here, in contrast to the enhancement observed in A549-ACE2 and HeLa-ACE2 cells, rapamycin, everolimus, and temsirolimus inhibited Spike-mediated infection in Calu-3 cells while ridaforolimus did not **(Supplemental Figure 4A)**. Furthermore, rapamycin, everolimus, and temsirolimus reduced IFITM3 protein in this cell line, but ridaforolimus had a negligible effect **(Supplemental Figure 4B)**. These results support that the effect of rapalog treatment on Spike-mediated infection is explained by their ability to induce the degradation of IFITM proteins, which inhibit SARS-CoV-2 infection in most contexts but enhance SARS-CoV-2 infection in Calu-3 cells for unknown reasons.

### Rapalogs differentially activate a lysosomal degradation pathway orchestrated by TFEB

Since rapamycin and rapalogs are known to inhibit mTOR signaling by binding both mTOR and FKBP12 (and other FKBP members), we sought to determine whether mTOR binding and its inhibition are required for rapalog-mediated enhancement of SARS-CoV-2 infection. To that end, we tested the effect of tacrolimus (also known as FK506), a macrolide immunosuppressant that is chemically related to rapalogs but does not bind nor inhibit mTOR. Instead, tacrolimus forms a ternary complex with FKBP12 and calcineurin to inhibit the signaling properties of the latter (50). In HeLa-ACE2 cells, a four-hour treatment of 20 μM tacrolimus did not reduce levels of IFITM2/3 **(Supplemental Figure 5A)**, nor did it boost HIV-CoV-2 infection **(Supplemental Figure 5B)**. These results suggest that FKBP12 binding is not sufficient for drug-mediated enhancement of SARS-CoV-2 infection. They also suggest that the extent to which mTOR is inhibited may explain the differential degree to which infection is impacted by the immunosuppressants examined in this study. Therefore, we surveyed the phosphorylation status of TFEB, a transcription factor that controls lysosome biogenesis and degradative processes carried out by lysosomes (51). mTOR phosphorylates TFEB at serine 211 (S211), which promotes its sequestration in the cell cytoplasm and decreases its translocation into the nucleus (51–53). Furthermore, this phosphorylation event was previously shown to be sensitive to inhibition by rapamycin and temsirolimus (52, 54). We found that rapamycin, everolimus, and temsirolimus significantly reduced S211 phosphorylation of endogenous TFEB in A549-ACE2 cells while ridaforolimus did so to a lesser extent **(Figure 6A-B)**. Furthermore, we measured the subcellular distribution of TFEB-GFP in HeLa-ACE2 treated with different compounds and found that rapamycin, everolimus, and temsirolimus induced a significantly greater accumulation of TFEB-GFP in the nucleus **(Figure 6C-D)**. Therefore, nuclear translocation of TFEB is associated with IFITM2/3 degradation and increased cellular susceptibility to SARS-CoV-2 Spike-mediated infection.

**Figure 6:**
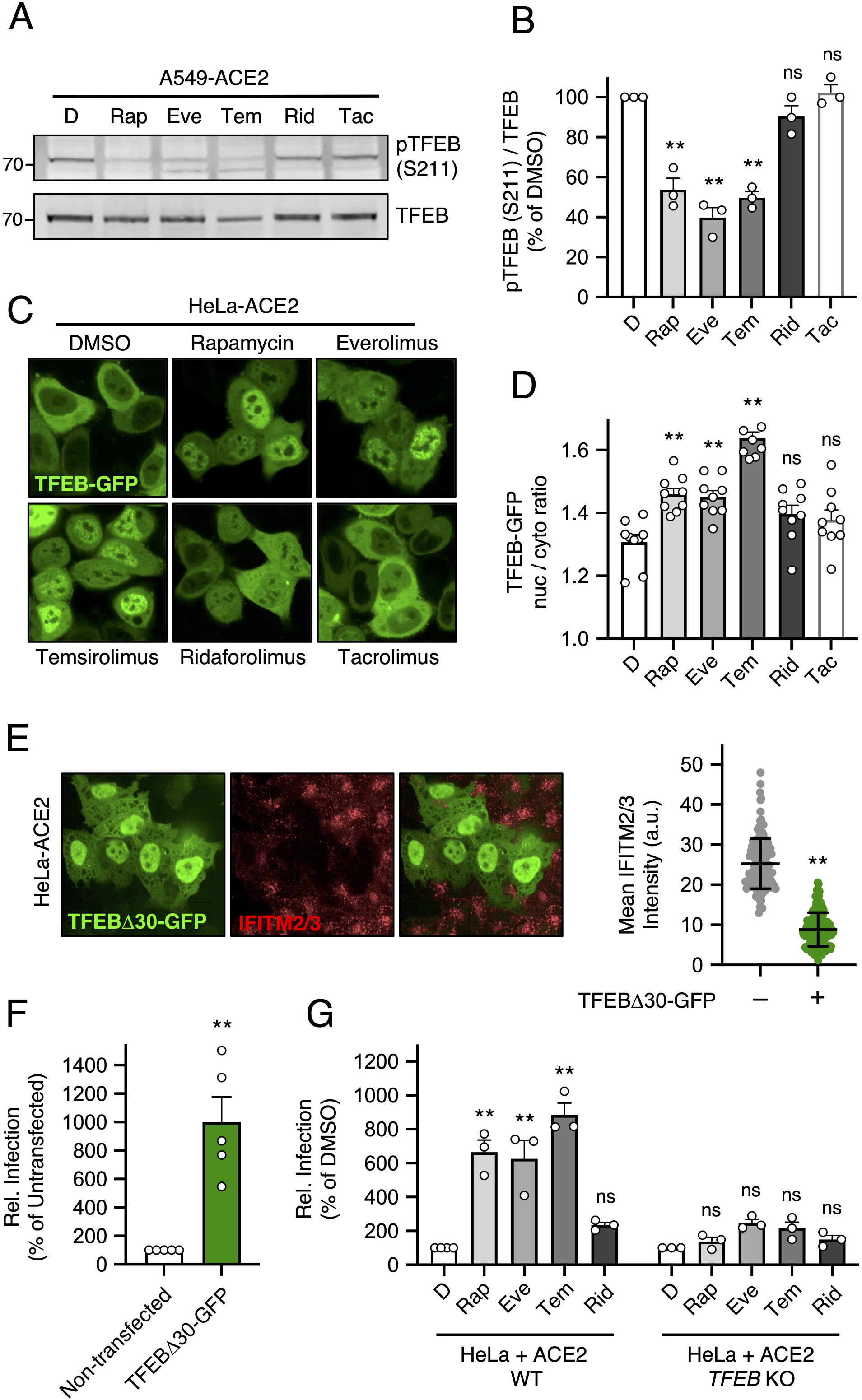
Nuclear TFEB triggers IFITM2/3 turnover, promotes Spike-mediated infection, and is required for enhancement of infection by rapalogs. (A) A549-ACE2 were treated with 20 µM Rap, Eve, Tem, Rid, tacrolimus (Tac), or DMSO for 4 hours and whole cell lysates were subjected to SDS-PAGE and Western blot analysis with anti-TFEB and anti-pTFEB (S211). (B) pTFEB (S211) levels were divided by total TFEB levels and summarized as an average of 3 experiments. (C) HeLa-ACE2 were transfected with TFEB-GFP for 24 hours, treated with Rap, Eve, Tem, Rid or Tac for 4 hours, stained with DAPI and CellMask (not shown), and imaged by high-content microscopy. Representative images are shown. (D) Ratio of nuclear to cytoplasmic TFEB-GFP was calculated in individual cells and average ratios derived from 9 separate fields of view (each containing 20-40 cells) are shown. (E) HeLa-ACE2 were transfected with 0.5 µg TFEBΔ30-GFP for 24 hours, fixed, stained with anti-IFITM2/3, and imaged by high-content microscopy (representative field on left). Average intensity of IFITM2/3 levels in 150 GFP-negative and 150 GFP-positive cells were grouped from two transfections (right). (F) HeLa-ACE2 were transfected (or not) with 0.5 µg TFEBΔ30-GFP, for 24 hours and HIV-CoV-2 (100 ng p24 equivalent) was added. Infection was measured by luciferase at 48 hours post-infection. Luciferase units were normalized to 100 in the non-transfected condition. (G) HeLa WT or *TFEB* KO were transfected with 0.3 µg pcDNA3.1-hACE2 for 24 hours and treated with 20 µM rapalogs/DMSO for 4 hours. HIV-CoV-2 (100 ng p24 equivalent) was added and luciferase activity measured at 48 hours post-infection. Luciferase units were normalized to 100 in the non-transfected condition. Means and standard error were calculated from 3 (A), 5 (F), and 3 (G) experiments. Statistical analysis was performed with one-way ANOVA or student’s T test (E and F) and asterisks indicate significant difference from DMSO or non-transfected conditions. *, p < 0.05; **, p < 0.01. Rel.; relative. A.u.; arbitrary units.

We confirmed that 20 μM ridaforolimus did not inhibit S211 phosphorylation of TFEB in HeLa-ACE2 cells, while the same concentration of temsirolimus did **(Supplemental Figure 6A-B)**. To better understand why ridaforolimus displayed less activity with regards to enhancing SARS-CoV-2 infection and inhibiting TFEB phosphorylation, we treated cells with increasing concentrations of ridaforolimus. Interestingly, we found that 30 μM ridaforolimus boosted infection to a similar extent as 20 μM temsirolimus, and 50 μM ridaforolimus boosted even further **(Supplemental Figure 6C)**. Further cementing the link between infection enhancement and nuclear translocation of TFEB, we found that elevated concentrations of ridaforolimus which resulted in increased infection were also sufficient to inhibit TFEB phosphorylation **(Supplemental Figure 6D)**. These findings indicate that, compared to other rapalogs, ridaforolimus is a less potent inhibitor of mTOR-mediated phosphorylation of TFEB, which may have important implications for the clinical use of ridaforolimus as an mTOR inhibitor in humans.

Consistent with a direct relationship between TFEB activation, IFITM2/3 turnover, and Spike-mediated cell entry, we found that ectopic expression of a constitutively active form of TFEB lacking the first 30 amino-terminal residues (51) was sufficient to trigger IFITM2/3 loss from cells **(Figure 6E)** and sufficient to increase susceptibility to HIV-CoV-2 infection **(Figure 6F)**. By combining transfection of the constitutively active form of TFEB with temsirolimus treatment, we found that IFITM2/3 levels were strongly suppressed irrespective of whether TFEB was detected or not. This confirms that TFEB and rapalogs are functionally redundant and operate in the same pathway to negatively regulate IFITM2/3 levels (**Supplemental Figure 7A**). Finally, we took advantage of TFEB-deficient cells to formally address the role that TFEB activation plays during rapalog-mediated enhancement of infection **(Supplemental Figure 7B)**. While rapamycin, everolimus, and temsirolimus significantly boosted HIV-CoV-2 infection in HeLa WT cells transfected with ACE2, no significant enhancement was observed in HeLa *TFEB* KO cells **(Figure 6G)**. In summary, our results employing functionally divergent rapalogs reveal a previously unrecognized immunoregulatory role played by the mTOR-TFEB-lysosome axis that affects the cell entry of SARS-CoV-2 and other viruses.

### Rapamycin enhances SARS-CoV-2 replication in primary human nasal epithelia and promotes viral disease in animal models

Our findings from SARS-CoV-2 and pseudovirus infection of human cells demonstrate that rapamycin, everolimus, and temsirolimus can suppress intrinsic immunity at the post-translational level, while ridaforolimus exhibits decreased potency in this regard. However, whether these compounds are functionally divergent when administered in vivo was unclear. To closely approximate the conditions under which SARS-CoV-2 infects and replicates within the human respiratory tract, we tested how rapamycin or ridaforolimus impacted SARS-CoV-2 replication in primary human nasal epithelial cells cultured at the liquid-air interface, a tissue model that recapitulates the 3D physiology of the upper airway. Measurement of viral *ORF1a* RNA by RT-qPCR was used to assess levels of viral transcripts at 24 and 48 hours post-infection, while *IL6* and *IFNB1* RNA were measured to assess the concomitant induction of cytokines. Levels of *ORF1a* significantly increased from 24 to 48 hours post-infection, suggesting that these cells support virus replication (**Figure 7A**). Furthermore, we found that rapamycin significantly enhanced virus replication (400-fold) at 48 hours post-infection, while ridaforolimus did not (**Figure 7A**). Consistent with enhanced virus replication in those cells, *IL6* and *IFNB1* transcripts were significantly elevated by rapamycin (**Figure 7B-C**). However, since rapamycin elevated viral *ORF1a* by 400-fold but only increased cellular *IL6* and *IFNB1* by 2.5-fold or less, these results suggest that rapamycin increases cellular susceptibility to SARS-CoV-2 infection while limiting inflammatory cytokine induction in response to infection.

**Figure 7:**
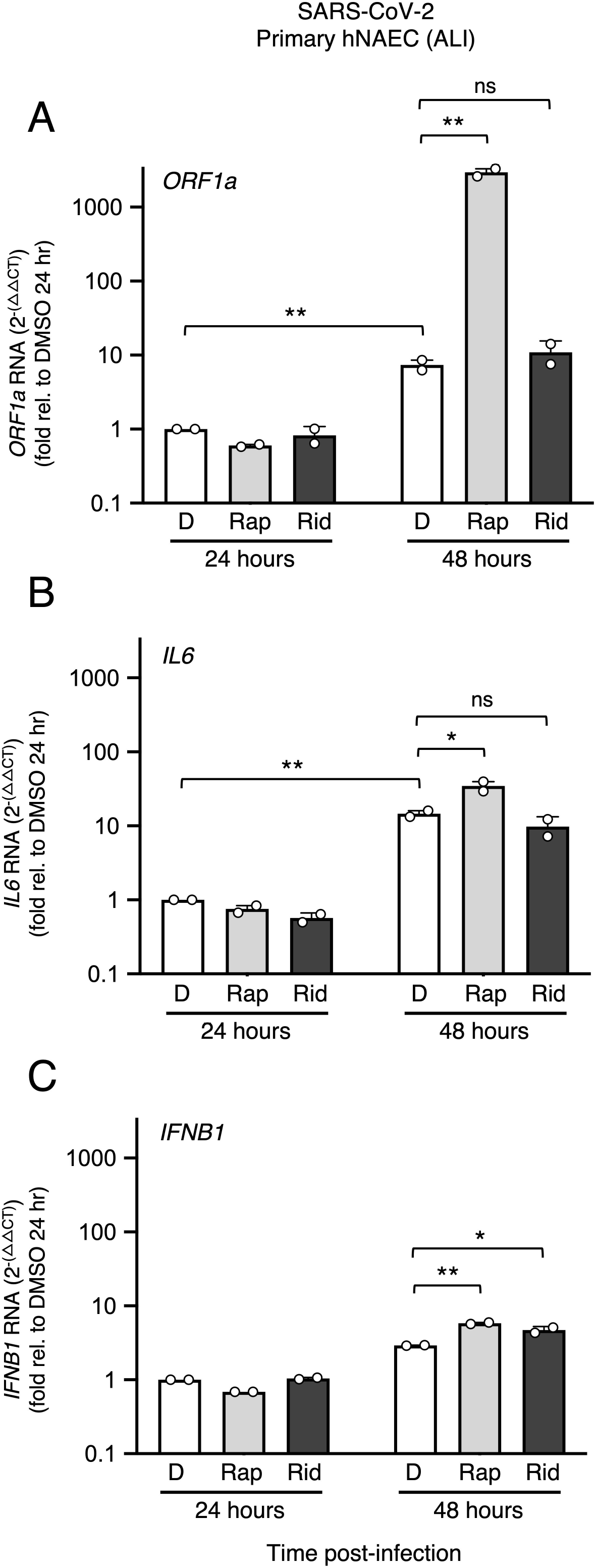
Rapamycin increases susceptibility of primary human nasal epithelial cells to SARS-CoV-2 infection while limiting pro-inflammatory cytokine induction. Primary human nasal epithelial cells (hNAEC) pooled from 12 donors were cultured at the liquid-air interface for 30-60 days were infected with 5^10^5^ plaque forming units (pfu) SARS-CoV-2 (WA1). At 24 hours and 48 hours post-infection, Trizol was added to cells and total RNA extraction was performed. RT-qPCR was performed using primers and probes specific to viral *ORF1a* (A), cellular *IL6* (B), and cellular *IFNB1* (C). Means and standard error were calculated from 2 experiments (infection of pooled cells from 12 human donors was performed in duplicate). Relative RNA levels are presented Comparative CT method with beta actin (*ACTB*) serving as an endogenous control. RNA levels present in the DMSO condition at 24 hours were normalized to 1. *ORF1a* was not detected in non-infected cells. Statistical analysis was performed using one way ANOVA. *, p < 0.05; **, p < 0.01. ns; not significant. rel.; relative.

Based on these findings, we next tested how intraperitoneal injection of rapamycin or ridaforolimus impacted SARS-CoV-2 replication and disease course in naive hamsters **(Figure 8A)**. Hamsters are a permissive model for SARS-CoV-2 because hamster ACE2 is sufficiently similar to human ACE2 to support productive infection. Furthermore, hamsters exhibit severe disease characterized by lung pathology when high viral loads are achieved (55). Eight hamsters were randomly allocated to each treatment group (rapamycin, ridaforolimus, or DMSO) and all received an intraperitoneal injection (3 mg/kg) 4 hours prior to intranasal inoculation with SARS-CoV-2 WA1. Furthermore, half of the hamsters in each group received a second injection on day 2 post-infection. As an indicator of viral disease, we tracked weight loss for 10 days, or less if the hamster met requirements for euthanasia (loss of 20% or more of its body weight or signs of respiratory distress such as agonal breathing). We observed that hamsters receiving two injections did not exhibit significantly different rates of weight loss compared to those receiving a single injection **(Supplemental Figure 8A)**. As a result, we consolidated hamsters into three groups of eight according to receipt of rapamycin, ridaforolimus, or DMSO. In addition to monitoring weight and breathing over the course of infection, disease scores (referred to as ‘COVID scores’) were generated daily for each hamster. Scoring reflected the extent of coat ruffling, hunched posture, lethargic state, and weight loss, and mean scores were compiled for each group. In agreement with the increased occurrence of morbidity necessitating euthanasia (**Figure 8B**), disease scores were higher on average for rapamycin- and ridaforolimus-treated hamsters relative to DMSO (**Supplemental Figure 8B and Supplemental Table 1**). Between days 6 and 8 post-infection, one (1/8) of the hamsters treated with DMSO exhibited severe morbidity necessitating euthanasia, while seven (7/8) of the hamsters treated with rapamycin did **(Figure 8B-C)**. Meanwhile, four (4/8) of the hamsters treated with ridaforolimus met requirements for euthanasia. Survivors in all three groups recovered weight after day 7 post-infection and infectious virus was not detected from the lungs of these hamsters at day 10 (**Figure 8D**). Overall, hamsters treated with rapamycin exhibited significantly reduced survival compared to the DMSO group, while survival of ridaforolimus-treated animals was decreased but did not differ significantly **(Figure 8C)**.

**Figure 8:**
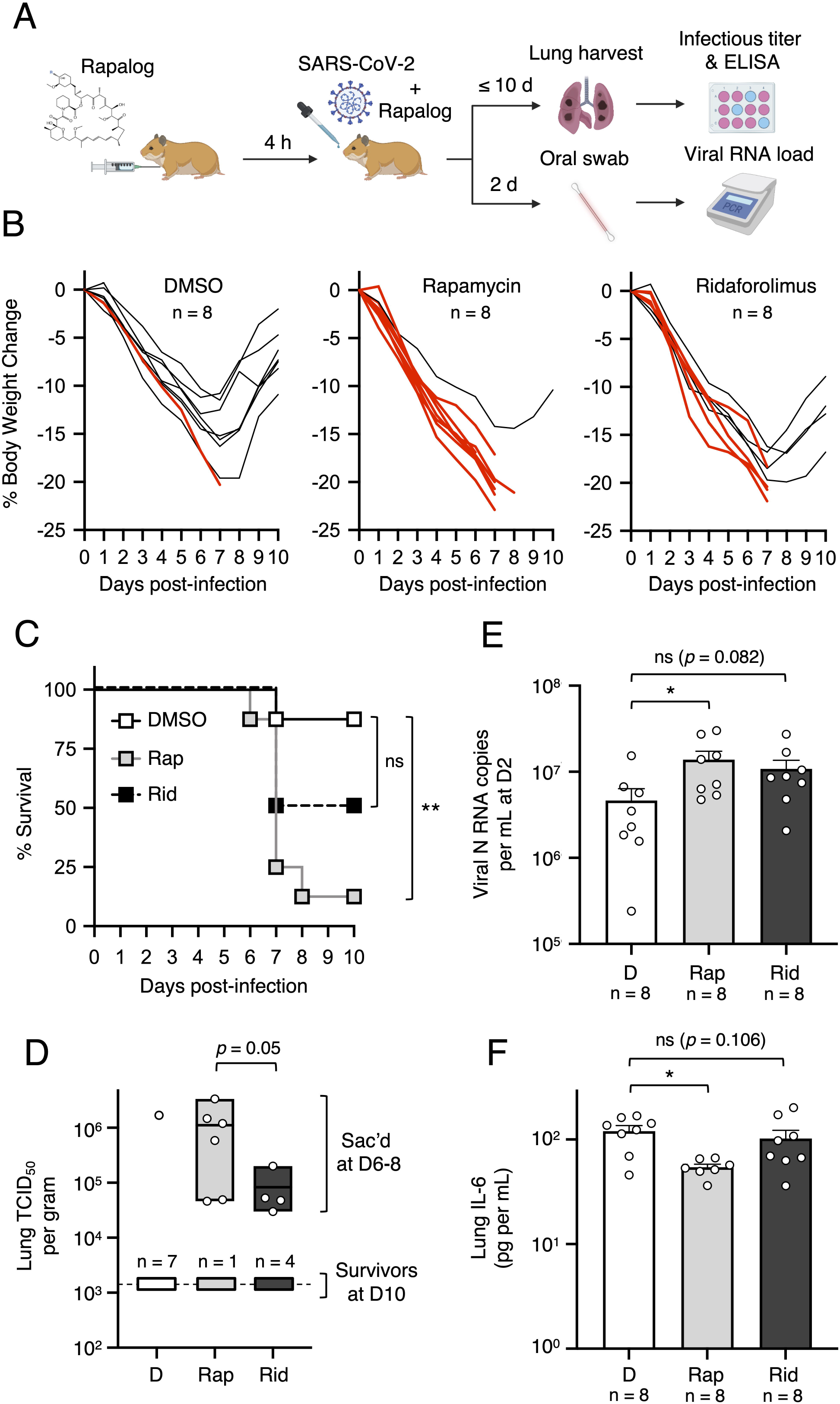
Rapamycin injection into hamsters intensifies viral disease during SARS-CoV-2 infection. (A) Golden Syrian hamsters were injected intraperitoneally with 3 mg/kg Rap, Rid, or equivalent amounts of DMSO (4 animals per group). Four hours later, hamsters were infected intranasally with 6 × 10^3^ plaque forming units of SARS-CoV-2. At 2 days post-infection, half of the animals received a second injection of Rap, Rid, or DMSO. Oral swabs were taken and used for measurement of oral viral RNA load by qPCR. At 10 days post-infection (or earlier, if more than 20% weight loss or agonal breathing was detected), hamsters were euthanized, and lungs were harvested for determination of infectious virus titer by TCID_50_ assay and IL-6 ELISA. (B) Individual body weight trajectories for each treatment group are plotted by day post-infection. Red lines indicate animals that required euthanasia for humane endpoints (more than 20% weight loss or agonal breathing). (C) Kaplan-Meier survival curves were generated according to the dates of euthanasia (or in one case, when an animal was found dead). (D) Infectious virus titers in lungs were determined by TCID_50_ in Vero-TMPRSS2 cells. Data is depicted as floating bars (minimum, maximum, and mean shown). (E) Viral RNA copy number was determined by qPCR from oral swab at 2 days post-infection. Data is depicted as box and whiskers plots. (F) IL-6 protein levels in lungs were determined using a hamster IL-6 ELISA kit. Statistical analysis in (C) was performed by comparing survival curves between Rap and DMSO or Rid and DMSO using the Log-rank (Mantel-Cox) test. Statistical analysis in (D) was performed by comparing all individuals (survivors and euthanized) in the Rap and Rid groups using the Mann-Whitney test. Statistical analysis in (E) and (F) was performed by one way ANOVA. Illustration created with BioRender.com. *, p < 0.05; **, p < 0.01. ns; not significant.

Lungs were harvested from infected hamsters following euthanasia either at the end of the experiment (for survivors) or earlier (for hamsters exhibiting morbidity and necessitating humane euthanasia). In contrast, the lungs of hamsters euthanized due to morbidity exhibited high infectious virus titers, suggesting that morbidity was caused by viral pathogenesis (the lungs of one hamster treated with rapamycin were not examined because it was found dead following infection) **(Figure 8D)**. In general, hamsters treated with rapamycin exhibited significantly higher infectious virus titers in lungs than those treated with ridaforolimus **(Figure 8D)**. In addition, early SARS-CoV-2 replication was measured by quantitative PCR from oral swabs. We found that hamsters injected with rapamycin exhibited significantly higher viral RNA levels in the oral cavity at day 2 post-infection compared to animals injected with DMSO **(Figure 8E)**. In contrast, viral RNA levels in hamsters injected with ridaforolimus were elevated relative to the DMSO group, but they did not differ significantly. Consistent with the known inhibitory effects of rapamycin on cytokine signaling (29), we detected significantly less IL-6 protein in lungs of hamsters treated with rapamycin, while ridaforolimus did not cause a reduction in IL-6 (**Figure 8F**). Overall, these results demonstrate that rapamycin administration increases host susceptibility to SARS-CoV-2 infection and significantly increases virus-induced morbidity in a manner that is not associated with an enhanced pro-inflammatory state.

These conclusions were supported by histopathological analysis of lungs, which indicated that lung damage was observed in all infected hamsters, especially those that needed to be humanely euthanized. All hamsters, regardless of treatment group, exhibited signs of lung hyperplasia and mixed or mononuclear inflammation, while some hamsters exhibited lung edema, hypertrophy, fibrosis, or syncytial cell formation. Hamsters requiring euthanasia, regardless of treatment group, showed the additional signs of moderate to severe lung hemorrhage, while minor hemorrhaging was apparent in only two hamsters that survived until day 10 post-infection (**Supplemental Table 1 and Appendix**). Since the highest viral loads in lungs were observed in morbid hamsters **(Figure 8D)**, lung dysfunction (acute respiratory distress syndrome) caused by virus replication is the likely cause of morbidity in hamsters. This is further supported by instances of agonal breathing in some of the infected hamsters, which necessitated euthanasia **(Supplemental Table 1)**.

Rapamycin was previously shown to promote morbidity of Influenza A infection in mice (36, 56). Moreover, we previously found that murine IFITM3 is sensitive to depletion by rapamycin (38). To determine whether rapamycin promotes host susceptibility to SARS-CoV-2 infection in mice, we injected C57BL/6 mice with rapamycin or DMSO prior to and after challenge with mouse-adapted (MA) SARS-CoV-2 **(Figure 9A)**. In this model, significant weight loss was not observed for up to five days following infection **(Supplemental Figure 8C)**. Lungs from mice in both groups were harvested uniformly on day 2 post-infection, and we found that virus titers were significantly increased (144-fold) in rapamycin-treated mice compared to DMSO-treated mice **(Figure 9B)**. As observed in hamsters, IL-6 levels were significantly reduced in lungs from rapamycin-treated mice despite enhanced virus titers **(Figure 9C)**. Furthermore, murine IFITM3 protein levels were reduced in the lungs of mice injected with rapamycin compared to levels found in DMSO-treated mice (**Figure 9D**). Together, these findings support the conclusion that rapamycin downmodulates cell-intrinsic barriers to SARS-CoV-2 infection in vivo, and as a result, enhances virus replication and viral disease.

**Figure 9:**
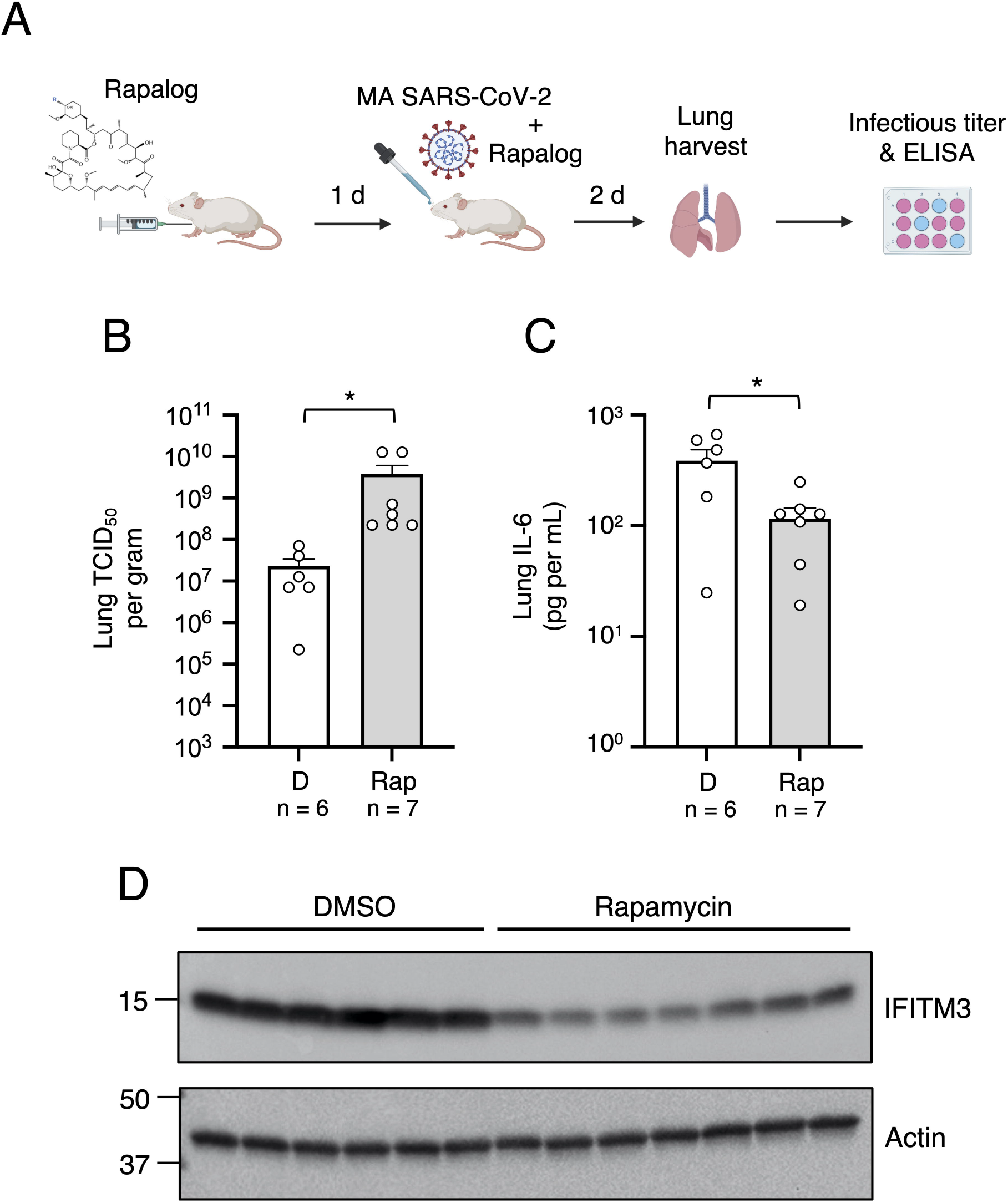
Rapamycin injection into mice downmodulates IFITM3 in lungs and boosts MA SARS-CoV-2 titers. (A) C57BL/6 mice were injected with 3 mg/kg of Rap or an equivalent amount of DMSO (6 or 7 mice per group, respectively). The following day, mice were infected intranasally with 6 × 10^4^ TCID_50_ mouse-adapted (MA) SARS-CoV-2. Mice received second and third injections of Rap or DMSO on the day of infection and on day 1 post-infection. (B) Lungs were harvested from infected mice upon euthanasia at day 2 post-infection and infectious viral loads were determined by TCID_50_ (B) and IL-6 protein was measured by a mouse IL-6 ELISA kit (C). Geometric mean TCID_50_ per gram was calculated per treatment group. Statistical analysis was performed with Mann-Whitney test and asterisks indicate significant difference from DMSO. *, p < 0.05; **, p < 0.01. (D) Lung homogenates (3 µg) from mice injected with Rap or DMSO were subjected to SDS-PAGE and Western blot analysis. Immunoblotting was performed with anti-Fragilis/IFITM3 (ab15592) and anti-actin. Illustration created with BioRender.com.

## Discussion

By assessing their impact on infection at the single-cell and whole-organism level, we draw attention to an immunosuppressive property of rapamycin and some rapalogs that acts on cell-intrinsic immunity and increases cellular susceptibility to infection by SARS-CoV-2 and likely other pathogenic viruses. Side effects of rapalog use in humans, including increased risk of respiratory tract infections, are regularly attributed to immunosuppression of adaptive immunity (57). Indeed, rapalogs have been used to mitigate systemic immunopathology caused by T-cell responses, and this is one reason why they are being tested for therapeutic benefit in COVID-19 patients. However, since rapamycin was injected into immunologically naive hosts prior to and soon after virus challenge and followed for no more than 10 days, it is unlikely that rapalogs modulated adaptive immunity against SARS-CoV-2 in our experiments. While immunomodulation of adaptive immunity by rapalogs may provide benefit for patients already suffering from COVID-19, pre-existing rapalog use may enhance host susceptibility to infection and disease by counteracting cell-intrinsic immunity.

The injection dose of rapamycin or ridaforolimus (3 mg/kg) that we administered to hamsters and mice, when adjusted for body surface area and an average human weight of 60 kg (58), equates to approximately 15 mg per human. This figure is similar to those administered to humans in clinical settings, such as the use of rapamycin for the treatment of glioblastoma (up to 10 mg daily for multiple days), the use of temsirolimus for the treatment of renal cell carcinoma (25 mg once weekly), or the use of everolimus for the treatment of tuberous sclerosis (TS), a genetic disorder resulting in hyperactivation of mTOR (10 mg daily, continuously) (23, 59–61). Interestingly, a case report detailed the deaths of two TS patients (a father and daughter) who, despite discontinuing everolimus upon detection of SARS-CoV-2 infection, died from severe COVID-19 in late 2020 (61). Our findings detailing the suppression of cell-intrinsic immunity by rapalogs raise the possibility that their use may predispose individuals to SARS-CoV-2 infection and severe forms of COVID-19. More generally, they provide new insight into how rapamycin and rapalogs may elicit unintended immunocompromised states and increase human susceptibility to multiple virus infections.

By leveraging the differential functional properties of rapalogs, we reveal how the mTOR-TFEB-lysosome axis impacts intrinsic resistance to SARS-CoV-2 infection. Specifically, rapamycin and select rapalogs (everolimus and temsirolimus) promote infection at the stage of cell entry, and this is functionally linked to nuclear accumulation of TFEB and the lysosomal degradation of IFITM proteins by endolysosomal microautophagy **(Figure 10)**. While mTOR phosphorylates TFEB at S211 to promote the sequestration of TFEB in the cytoplasm, the phosphatase calcineurin dephosphorylates TFEB at this position to promote nuclear translocation (62). Therefore, the extent to which different rapalogs promote nuclear TFEB accumulation may be a consequence of differential mTOR inhibition and/or differential calcineurin activation. Calcineurin is activated by calcium release through the lysosomal calcium channel TRPML1 (also known as mucolipin-1) (62), and interestingly, it was shown that rapamycin and temsirolimus, but not ridaforolimus, promote calcium release by TRPML1 (54). Therefore, it is worth examining whether TRPML1 or related lysosomal calcium channels are required for the effects of rapalogs on virus infection. Overall, our findings reveal a previously unrecognized mechanism by which TFEB promotes virus infections—inhibition of cell-intrinsic defenses restricting virus entry. We show that nuclear TFEB induces the degradation of IFITM proteins, but it may also trigger the loss or relocalization of other antiviral factors that remain to be uncovered. Furthermore, TFEB-mediated induction of dependency factors, such as cathepsin L, is likely to partially contribute to the overall impact of rapalogs on SARS-CoV-2 infection. Overall, this work identifies TFEB as a therapeutic target, and inhibitors that limit levels of nuclear TFEB could be mobilized for broad-spectrum antiviral activity.

**Figure 10:**
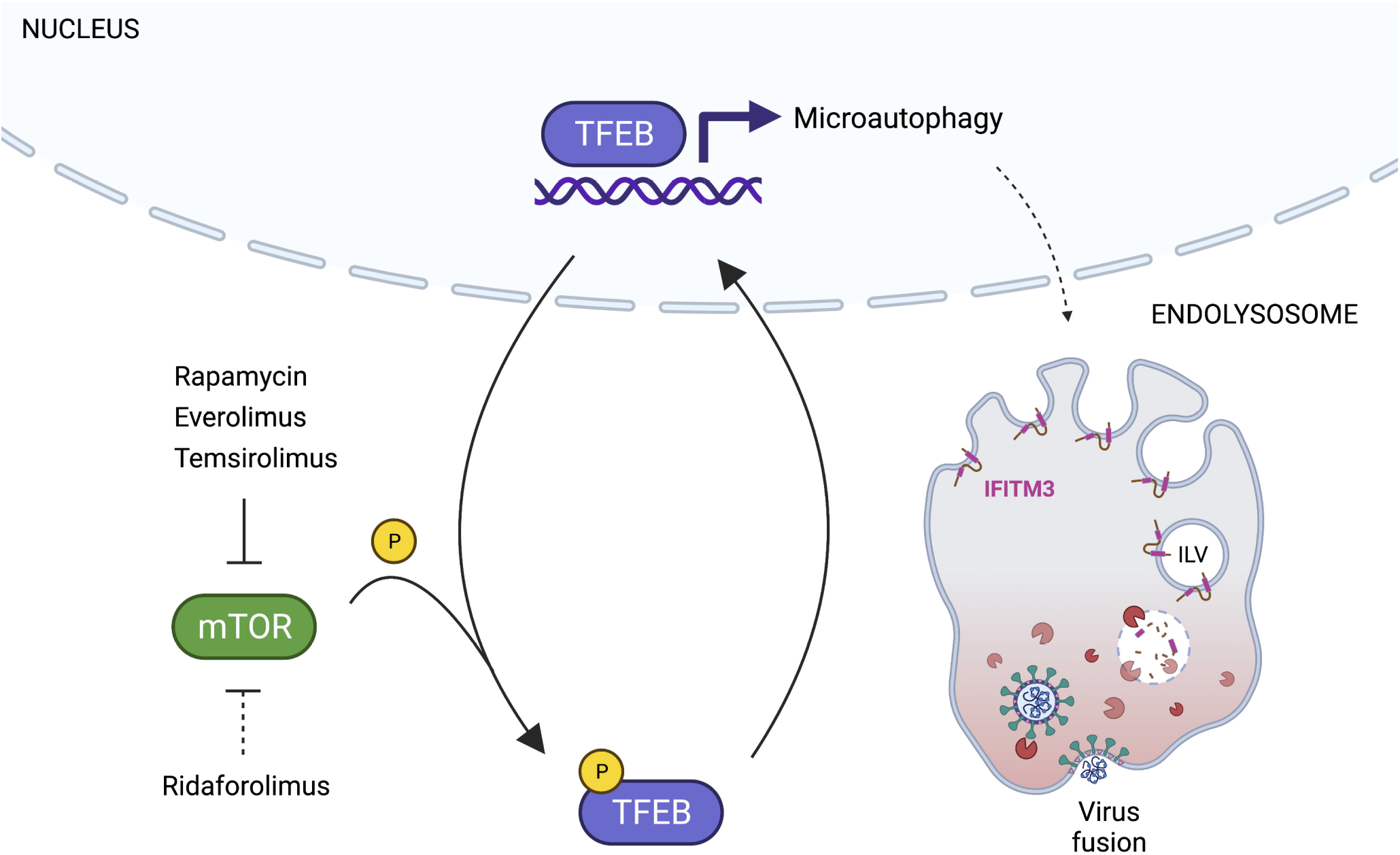
Model for rapalog-mediated enhancement of SARS-CoV-2 infection. Rapamycin, everolimus, and temsirolimus potently inhibit the phosphorylation of TFEB by mTOR, while ridaforolimus is a less potent inhibitor. As a result, TFEB translocates into the nucleus and induces genes functioning in lysosomal activities, including autophagy-related pathways. Nuclear TFEB triggers a microautophagy pathway that results in accelerated degradation of membrane proteins IFITM2 and IFITM3. Loss of IFITM2/3 promotes SARS-CoV-2 entry into cells by facilitating fusion between viral membranes and cellular membranes. Illustration created with BioRender.com.

We previously demonstrated that treatment of cells with micromolar quantities of rapamycin induced the lysosomal degradation of IFITM2/3 via a pathway that is independent of macroautophagy yet dependent upon endosomal complexes required for transport (ESCRT)-mediated sorting of IFITM2/3 into intraluminal vesicles of late endosomes/MVB (38). This MVB-mediated degradation pathway is also referred to as microautophagy, which occurs directly on endosomal or lysosomal membranes and involves membrane invagination (63). In both yeast and mammalian cells, microautophagy is characterized by ESCRT-dependent sorting of endolysosomal membrane proteins into intraluminal vesicles followed by their degradation by lysosomal hydrolases (64). While microautophagy selectively targets ubiquitinated endolysosomal membrane proteins, cytosolic proteins can also be non-selectively internalized into intraluminal vesicles and degraded (65, 66). Interestingly, microautophagy is known to be regulated by mTOR (67, 68), and mTOR inhibition triggers a ubiquitin- and ESCRT-dependent turnover of vacuolar (lysosomal) membrane proteins in yeast (69, 70). Overall, our findings suggest that select rapalogs induce a rapid, TFEB-dependent, endolysosomal membrane remodeling program known as microautophagy, and IFITM proteins are among the client proteins subjected to this pathway. The full cast of cellular factors that orchestrate this selective degradation program in mammalian cells and the other client proteins subjected to it will need to be worked out. Interestingly, the E3 ubiquitin ligase NEDD4 was previously shown to ubiquitinate IFITM2 and IFITM3 and to induce their lysosomal degradation in mammalian cells (71, 72), while Rsp5, the yeast ortholog of NEDD4, was shown to ubiquitinate vacuolar proteins turned over by microautophagy in yeast (73). Therefore, rapamycin and select rapalogs may upregulate NEDD4 function, resulting in selective degradation of a subset of the cellular proteome that includes IFITM proteins. Indeed, NEDD4 and the related NEDD4L are among the known target genes regulated by TFEB (74).

The relationship between IFITM proteins and human coronaviruses is complex. It was previously shown that IFITM3 facilitates replication of the seasonal coronavirus hCoV-OC43 (75), while we and others recently showed that SARS-CoV-1 and SARS-CoV-2 infection is inhibited by ectopic and endogenous IFITM1, IFITM2, and IFITM3 from mice and humans (47, 76–79). Intriguingly, mutants of human IFITM3 that lack the capacity to internalize into endosomes lost antiviral activity and promoted SARS-CoV-2 and MERS-CoV infection, revealing that IFITM3 can either inhibit or enhance infection depending on its subcellular localization (47, 80). Furthermore, one study reported that endogenous human IFITM proteins promoted infection by SARS-CoV-2 in certain human tissues, possibly by acting as interaction partners and docking platforms for viral Spike (81). Overall, the net effect of human IFITM proteins on SARS-CoV-2 infection in vivo remains unclear. However, the impact of rapamycin in our experimental SARS-CoV-2 infections of hamsters and mice suggests that rapamycin-mediated loss of IFITM proteins favors virus infection and viral disease, consistent with IFITM proteins performing antiviral roles against SARS-CoV-2 in those species. Accordingly, it was recently demonstrated that mouse IFITM3 protects mice from viral pathogenesis following MA SARS-CoV-2 infection (82).

Other lines of evidence support an antiviral role for IFITM proteins during SARS-CoV-2 infection in humans. While SARS-CoV-2 infection has been shown to cause deficiencies in interferon synthesis and interferon response pathways, administration of type I interferon in vivo promotes SARS-CoV-2 clearance in hamsters and humans (83). Notably, IFITM3 is among the most highly induced genes in primary human lung epithelial cells exposed to SARS-CoV-2 (84, 85), and humans experiencing mild or moderative COVID-19 showed elevated induction of antiviral genes, including *IFITM1* and *IFITM3*, in airway epithelium compared to individuals suffering from more severe COVID-19 (86). Single nucleotide polymorphisms in human *IFITM3* known as ns12252 and rs34481144, which lead to IFITM3 loss-of-function, have been associated with severe outcomes following Influenza A virus infection as well as severe COVID-19 (87, 88). These data suggest that cell-intrinsic immunity in airways plays a role in restricting virus spread and constraining systemic pathology during infection. Therefore, downmodulation of IFITM proteins by select rapalogs may contribute to the immunocompromised state that these drugs are well known to elicit in humans. This possibility warrants the close examination of different rapalog regimens on respiratory virus acquisition and disease in humans.

## Methods

### Cell lines, cell culture, inhibitors, and cytokines

HEK293T (CRL-3216) and Calu-3 (HTB-55) cells were obtained from ATCC. HeLa-ACE2, HeLa-DPP4, and A549-ACE2 cell lines were produced by transducing cells with lentivirus packaging pWPI encoding ACE2 or DPP4 and selecting with blasticidin. HeLa IFITM1/2/3 Knockout (C5-9) cells were purchased from ATCC (CRL-3452). HeLa *TFEB* KO cells were kindly provided by Ramnik J. Xavier (Broad Institute) and were described in (89). Primary human small airway (lung) epithelial cells (HSAEC) were purchased from ATCC (PCS-301-010). The partially immortalized nasal epithelial cell line (UNCNN2TS) was kindly provided by Scott H. Randell (University of North Carolina School of Medicine). Vero E6 cells (NR-53726) were obtained from BEI Resources. Vero-TMPRSS2 cells were a kind gift from Shan-Lu Liu (The Ohio State University). All cells were cultured at 37°C with 5% CO_2_ in Dulbecco’s Modified Eagle Medium (DMEM) supplemented with 10% fetal bovine serum (HyClone, Cytiva), except for UNCNN2TS, which were cultured in EpiX Medium (Propagenix), and HSAEC, which were cultured with airway epithelial cell basal medium (ATCC, PCS-300-030) and the bronchial epithelial cell growth kit (ATCC, PCS-300-040). Primary human nasal airway epithelial cells (hNAEC) cultured at the air-liquid interface were obtained from Epithelix (EP02MP, MucilAir Pool of Donors) and cultured according to the provider’s instructions using MucilAir culture medium. Rapamycin (553211) was obtained from Sigma. Everolimus (S1120), temsirolimus (S1044), ridaforolimus (S5003), tacrolimus (S5003), and SAR405 (S7682) were obtained from Selleckchem. U18666A (U3633) and Bafilomycin A1 (SML1661) were obtained from Sigma. Type-I interferon (human recombinant interferon-beta_ser17_, NR-3085) was obtained from BEI Resources.

### Plasmids and RNA interference

pcDNA3.1 encoding human ACE2 was kindly provided by Thomas Gallagher (Loyola University). pcDNA3.1 encoding CoV-1 Spike or CoV-2 Spike (WA1) tagged with a C9 epitope on the C-terminus, or MERS Spike, was kindly provided by Thomas Gallagher (Loyola University). pcDNA3.1 encoding CoV-1 Spike or CoV-2 Spike (WA1) tagged with a FLAG epitope on the C-terminus was obtained from Michael Letko and Vincent Munster (NIAID). pcDNA3.1 encoding CoV-2 Omicron (BA.1) Spike tagged with a His epitope on the N-terminus was synthesized provided by Genscript. pMD2.G encoding VSV-G (12259) was obtained from Addgene (a generous gift from Didier Trono). pWPI was obtained from Addgene (12254) and human ACE2 or human TMPRSS2 was introduced by Gateway cloning (Gateway LR Clonase II Enzyme mix (11791020)) as per manufacturer’s instructions. pPolII encoding hemagglutinin (HA) or neuraminidase (NA) from Influenza A/Turkey/1/2005 (H5N1) were kindly provided by Richard Yi Tsun Kao (The University of Hong Kong). pCMV encoding HIV-1 Vpr fused to beta lactamase (pCMV4-BlaM-Vpr) was obtained from Addgene (21950). A plasmid encoding replication-incompetent HIV-1 lacking *env* and *vpr* and encoding luciferase (pNL4-3LucR-E-) was kindly provided by Vineet KewalRamani (National Cancer Institute). A plasmid encoding replication-incompetent HIV-1 lacking *env* (pNL4-3E-) was kindly provided by Olivier Schwartz (Institut Pasteur). pEGFP-N1-TFEB (38119) and pEGF-N1-Δ30TFEB (44445) were obtained from Addgene (a generous gift of Shawn M. Ferguson). pEGFP-2xFYVE (140047) was obtained from Addgene (a gift from Harald Stenmark). Silencer Select siRNA targeting IFITM3 (s195035) and a non-targeting control (No. 1) was obtained from Ambion. Cells were transfected with 20 nM siRNA using Opti-MEM (Gibco) and Lipofectamine RNAiMAX (Thermo Fisher).

### Virus and pseudovirus infections

SARS-CoV-2 isolate USA-WA1/2020 (MN985325.1) was provided by the Centers for Disease Control or by BEI Resources (NR-52281). Virus propagation was performed in Vero E6 cells. Mouse-adapted (MA) SARS-CoV-2 variant MA10 (in the USA-WA1/2020 backbone) (90) was obtained from BEI Resources (NR-55329). Virus propagation was performed in Vero E6 cells and subsequently in Vero-TMPRSS2 cells. Virus was sequenced to ensure lack of tissue culture adaptations, including furin cleavage site mutations. Virus titers were calculated by plaque assay performed in Vero E6 cells as follows: serial 10-fold dilutions were added to Vero E6 monolayers in 48-well plates for 1 hour at 37°C. Cells were overlayed with 1.5% carboxymethyl cellulose (Sigma) in modified Eagle’s medium containing 3% fetal bovine serum (Gibco), 1 mM L-glutamine, 50 units per mL penicillin and 50 µg per mL streptomycin. Three days post-infection, cells were fixed in 10% formalin and stained with crystal violet to visualize and count plaques as previously described (91). Titers were calculated as plaque forming units per mL and normalized as described in the figure captions. HIV-based pseudovirus was produced by transfecting HEK293T cells with 12 µg of pNL4-3LucR-E- and 4 µg of plasmid encoding viral glycoproteins (pcDNA3.1 Spike (CoV-1, CoV-2 WA1, CoV-2 Omicron/BA.1, or MERS-CoV), pMD2.G-VSV-G, or 2 µg of pPol1II-HA and 2 µg of pPol1II-NA) using TransIT-293 (Mirus). Virus supernatant was harvested 72 hours post-transfection and filtered through 0.22 µm filters. Pseudovirus titers were determined by p24 ELISA (XpressBio) and 100 ng p24 equivalent was added to target cells and incubated for 72 hours prior to lysis with Passive Lysis Buffer (Promega). Luciferase activity was measured using the Luciferase Assay System (Promega). VSV-based pseudovirus was produced as previously described (92). In brief, HEK293T cells were transfected with 2 µg pcDNA3.1 CoV-2 Spike using Lipofectamine2000 (Thermo Fisher). At 24 hours post-transfection, culture medium was removed from cells and 2 mL of VSV-luc/GFP + VSV-G (seed particles) was added. At 48 hours post-infection, virus supernatants were collected, clarified by centrifugation at 500xG for 5 mins, and stored. 50 µL of virus supernatants were added to target cells for a period of 24 hours prior to fixation with 4% paraformaldehyde (for measurements of GFP+ cells with flow cytometry). For infections with replication-competent SARS-CoV-2 (WA1) assessed by plaque assay, rapamycin, everolimus, temsirolimus, or ridaforolimus (20 µM) were used to pretreat cells for 4 hours and then drugs were washed away prior to addition of virus at a multiplicity of infection (MOI) of 0.1. DMSO (Sigma) was used as a vehicle control. At one-hour post-virus addition, cells were washed once with 1X PBS and overlayed with complete medium. Supernatants were harvested 24 hours later, and titers were determined on plaque assays performed in Vero E6 cells. For infections with replication-competent SARS-CoV-2 (WA1) assessed by RT-qPCR, primary hNAEC cultured at the liquid-air interface for 30-60 days were washed three times with PBS and treated with 20 µM rapamycin, ridaforolimus, or an equivalent volume of DMSO for 4 hours. Then 5^10^5^ plaque forming units were added to cells for 2 hours. Afterwards, inoculum and compound were removed, and the cells were washed three times with PBS. At 24- and 48-hours post-infection, Trizol was added to the cells and RNA extraction and RT-qPCR was performed. For single-round infections using HIV- or VSV-based pseudovirus, rapamycin, everolimus, temsirolimus, ridaforolimus, or tacrolimus (20 µM) were used to pretreat cells for 4 hours and were maintained for the duration of infection and until harvest of cells for luciferase assay or flow cytometry. DMSO (Sigma) was used as a vehicle control.

### FRET-based virus entry assay

HIV-based pseudovirus incorporating BlaM-Vpr and CoV-2 Spike was produced by transfecting HEK293T cells with pNL4-3E-(15 µg), pCMV4-BlaM-Vpr (5 µg), and pcDNA3.1 CoV-2 Spike (5 µg) using the calcium phosphate technique. Briefly, six million 293T cells were seeded in a T75 flask. Plasmid DNA was mixed with sterile H2O, CaCl2, and Tris-EDTA (TE) buffer, and the totality was combined with Hepes-buffered saline (HBS). The transfection volume was added dropwise, and cells were incubated at 37°C for 48 h. Supernatants were recovered and clarified by centrifugation, passed through a 0.45 µm filter, and stored. Titers were measured using an HIV-1 p24 ELISA kit (XpressBio). 50 ng p25 equivalent of virus was added to HeLa-ACE2 cells for 2 hours. Cells were washed and labeled with the CCF2-AM β-lactamase Loading Kit (Invitrogen) for 2 hours and analyzed for CCF2 cleavage by flow cytometry as described (93). Rapamycin, everolimus, temsirolimus, or ridaforolimus (20 µM) were used to pretreat cells for 4 hours prior to virus addition and were maintained for the duration of infection. DMSO (Sigma) was used as a vehicle control.

### Western blot, antibodies, and flow cytometry

Whole cell lysis was performed with RIPA buffer (Thermo Fisher) supplemented with Halt Protease Inhibitor EDTA-free (Thermo Fisher). Lysates were clarified by centrifugation and supernatants were collected and stored. Protein concentration was determined with the Protein Assay Kit II (Bio-Rad), and 10-15 µg of protein was loaded into 12% acrylamide Criterion XT Bis-Tris Precast Gels (Bio-Rad). Electrophoresis was performed with NuPage MES SDS Running Buffer (Invitrogen) and proteins were transferred to Amersham Protran Premium Nitrocellulose Membrane, pore size 0.20 µm (GE Healthcare). Membranes were blocked with Odyssey Blocking Buffer (Li-COR) and incubated with the following primary antibodies diluted in Odyssey Antibody Diluent (Li-COR): anti-IFITM1 (60074-1-Ig; Proteintech), anti-IFITM2 (66137-1-Ig; Proteintech), anti-IFITM3 (EPR5242, ab109429; Abcam), anti-Fragilis (ab15592; Abcam (detects murine IFITM3)), anti-IFITM2/3 (66081-1-Ig; Proteintech), anti-actin (C4, sc-47778; Santa Cruz Biotechnology), anti-hACE2 (ab15348; Abcam), anti-TFEB (4240S; Cell Signaling Technology), and anti-pTFEB (Ser211) (37681S; Cell Signaling Technology). Secondary antibodies conjugated to DyLight 800 or 680 (Li-Cor) and the Li-Cor Odyssey CLx imaging system were used to reveal specific protein detection. Images were analyzed (including signal quantification) and assembled using ImageStudioLite (Li-Cor). Cell viability was measured using LIVE/DEAD Red Dead Cell Stain Kit (Thermo Fisher). Cells were fixed and permeabilized with Cytofix/Cytoperm reagent (BD) for 20 minutes and washed in Perm/Wash buffer (BD). Flow cytometry was performed on an LSRFortessa (BD).

### Confocal fluorescence and immunofluorescence microscopy

HeLa-ACE2 cells were fixed with 4% paraformaldehyde, stained with anti-IFITM2/3 (66081-1-Ig; Proteintech), goat anti-mouse IgG Alexa Fluor 647 (A21235; Thermo Fisher) and DAPI (62248; Thermo Fisher), and imaged in a glass-bottom tissue culture plate with an Operetta CLS High-Content Analysis System (Perkin Elmer). For measurement of TFEB-GFP nuclear/cytoplasmic distribution, HeLa-ACE2 cells were transfected with pEGFP-N1-TFEB for 24 hours, fixed with 4% paraformaldehyde, stained with HCS CellMask Red Stain (H32712; Thermo Fisher) and DAPI, and imaged with an Operetta CLS. Using Harmony software (Perkin Elmer), nuclear/cytoplasmic ratios of TFEB-GFP were calculated in single cells as follows: cells were delineated by CellMask Red Stain, nuclei were delineated by DAPI, nuclear TFEB-GFP was designated as GFP overlapping with DAPI, and cytoplasmic TFEB-GFP was designated as total GFP signal minus nuclear TFEB-GFP. Average ratios were calculated from 20-30 cells per field, and the mean of averages from 10 fields was obtained (total of approximately 250 cells per condition). For measurement of IFITM2/3 levels in cells transfected with TFEBΔ30-GFP, HeLa-ACE2 cells were transfected with pEGF-N1-Δ30TFEB for 24 hours, fixed and permeabilized with BD Cytofix/Cytoperm (Fisher Scientific), stained with anti-IFITM2/3 and goat anti-mouse IgG Alexa Fluor 647, and imaged with an Operetta CLS. The IFITM2/3 fluorescence intensity within a single, medial Z section was measured in approximately 150 GFP-negative cells and 150 GFP-positive cells using the freehand selections tool in ImageJ.

### RT-qPCR of viral and cellular transcripts in infected primary human nasal epithelial cells

Cells lysted with Trizol were mixed with chloroform (Sigma) at a 5:1 (Trizol:chloroform) ratio. Mixed samples were mixed thoroughly and incubated at room temperature for 10 minutes, followed by centrifugation at 12000 x G for 5 minutes to allow separation of the aqueous and organic phases. Equal volumes of 70% ethanol were added to the aqueous phases, mixed thoroughly, and incubated at room temperature for 5 minutes. RNA purification was performed using the PureLink RNA Mini Kit (Invitrogen) according to manufacturer’s instructions. Purified RNA product was immediately used with the One-step PrimeScript RT-PCR Kit (Takara). Primers and probes were obtained from IDT. The primers and probes used to amplify and quantify *ORF1a* are as follows (5’-3’): ORF1a-F AGAAGATTGGTTAGATGATGATAGT; ORF1a-R TTCCATCTCTAATTGAGGTTGAACC; ORF1a-P FAM/TCCTCACTGCCGTCTTGTTGACCA/BHQ13. The primers and probes used to amplify and quantify *IL6* are as follows (5’-3’): IL6-F GCAGATGAGTACAAAAGTCCTGA; IL6-R TTCTGTGCCTGCAGCTTC; IL6-P 56-FAM/CAACCACAA/ZEN/ATGCCAGCCTGCT/31ABkFQ. The primers and probes used to amplify and quantify *IFNB1* are as follows (5’-3’): IFNB1-F GAAACTGAAGATCTCCTAGCCT; IFNB1-R GCCATCAGTCACTTAAACAGC; IFNB1-P 56-FAM/TGAAGCAAT/ZEN/TGTCCAGTCCCAGAGG/3IABkFQ. The primers and probes used to amplify and quantify *ACTB* are as follows (5’-3’): ACTB-F ACAGAGCCTCGCCTTTG; ACTB-R CCTTGCACATGCCGGAG; ACTB-P 56-FAM/TCATCCATG/ZEN/GTGAGCTGGCGG/31ABkFQ. Reaction mixtures of 20 µL (including 2.2 µL total RNA, 0.2 µM forward and reverse primers, and 0.1 µM probe) were subjected to reverse transcription (5 min at 45°C, followed by 10 s at 95°C) and 40 cycles of PCR (5 s at 95°C followed by 20 s at 60°C) in a CFX Opus 96 Real-Time PCR System (Bio-Rad). Results were analyzed by the Comparative CT Method (ΔΔCt Method). RNA levels for *ORF1a, IL6*, and *IFNB1* in each sample were normalized to *ACTB*.

### In vivo infections of hamsters and mice with SARS-CoV-2

Male Golden Syrian hamsters between the ages of 6-8 weeks were acclimated for 11 days following receipt. Hamsters received an intraperitoneal injection (500 µL) of rapamycin (HY-10219; MedChemExpress) or ridaforolimus (HY-50908; MedChemExpress) at 3 mg/kg or an equivalent amount of DMSO (8 hamsters per group). Four hours later, hamsters were challenged with 6 × 10^3^ plaque forming units of SARS-CoV-2 isolate USA-WA1/2020 (amplified on Calu-3 cells) through intranasal inoculation (50 µL in each nare). Half of the hamsters in each group received a second injection at day 2 post-infection. Clinical observations and weights were recorded daily up until day 10 post-infection. According to Institutional Animal Care and Use Committee human euthanasia criteria, hamsters were euthanized immediately if weight loss exceeded 20% or if agonal breathing was detected. Otherwise, hamsters were euthanized on day 10 post-infection. Oral swabs were collected on day 2 post-infection for measurement of viral RNA by quantitative PCR of the viral N (nucleocapsid) gene. Lungs were harvested following euthanasia (day 10 or earlier) and infectious viral load was determined by TCID_50_ assay in Vero-TMPRSS2 cells. Histopathologic analysis of hamster lungs was performed by Experimental Pathology Laboratories, Inc. At necropsy, the left lung lobe was collected and placed in 10% neutral buffered formalin and processed to hematoxylin and eosin stained slides and examined by a board-certified pathologist. Histopathologic findings are presented in Appendix and Supplemental Table 1. Findings were graded from one to five (increasing severity). Male C57BL/6 mice received an intraperitoneal injection of 3 mg/kg rapamycin (NC9362949; LC-Laboratories) or an equivalent amount of DMSO (7 and 6 mice per group, respectively). The following day, mice were challenged intranasally with 5 × 10^4^ TCID_50_ equivalent of MA10 SARS-CoV-2 (USA-WA1/2020 backbone). Mice received a second injection of rapamycin or DMSO on the day of infection and a third on day one post-infection. Mice were euthanized for lung harvest on day two post-infection. Infectious viral load was determined by TCID_50_ assay in Vero-TMPRSS2 cells. Following UV-inactivation of lung homogenates, IL-6 protein was detected by Hamster IL-6 Sandwich ELISA Kit (AssayGenie) or Mouse IL-6 Duoset Sandwich ELISA kit (R&D Systems) according to manufacturers’ instructions. Animal studies were conducted in compliance with all relevant local, state, and federal regulations and were approved by the Institutional Animal Care and Use Committee of Bioqual and of the Ohio State University.

### Statistics

The statistical tests performed in each figure are described in the accompanying figure legend. In general, the cutoff (alpha) for significance was 0.05 and two-tailed tests were always performed.

### Study approval

Animal studies were conducted in compliance with all relevant local, state, and federal regulations and were approved by the Institutional Animal Care and Use Committee of Bioqual and of the Ohio State University.

## Author Contributions

AAC and GS designed the research studies and wrote the manuscript. GS, AIC, TL, AK, SM, KKL, TD, AZ, AE, LZ, and SK conducted experiments, acquired data, and analyzed data. PAB provided reagents. JWY, SMB, JSY, and AAC obtained funding and supervised the experiments. All authors contributed to editing of the manuscript.

## Acknowledgements

We thank Michael Letko and Vincent Munster for providing VSV-luc/GFP seed particles and CoV and CoV-2 Spike plasmids, Thomas Gallagher for providing CoV, CoV-2, and MERS Spike plasmids, Alan Rein for facilitating lentiviral pseudovirus production, Scott H. Randell for providing UNCNN2TS, Ramnik J. Xavier for providing HeLa *TFEB* KO cells, Eric O. Freed for providing primary HSAEC, the Integrated Research Facility (NIAID) at Fort Detrick for providing *ORF1a* primers and probe, and the SARS-CoV-2 Virology Core (NIAID) for use of their dedicated BSL3 laboratory space.

