## Supplemental Figures for "Rapalogs downmodulate intrinsic immunity and promote cell entry of SARS-CoV-2"

Supplemental Figure 1

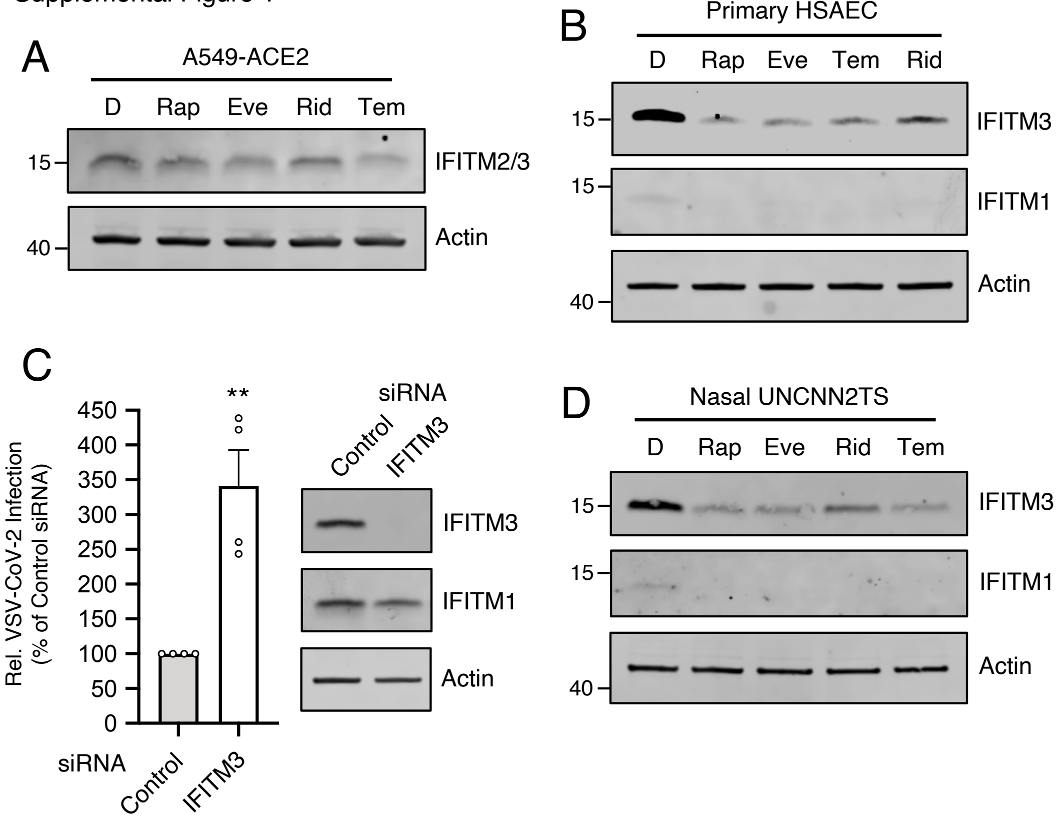


**Supplemental Figure 1:** (A) A549-ACE2 cells were treated with 20 μM Rap, Eve, Rid, Tem, or an equivalent volume of DMSO (in the absence of type-I interferon) for 4 hours and whole cell lysates were subjected to SDS-PAGE and Western blot analysis. Immunoblotting was performed with anti-IFITM2/3 and anti-actin. (B) Primary HSAEC were treated with 20 μM Rap, Eve, Tem, Rid, or an equivalent volume of DMSO for 4 hours and whole cell lysates were subjected to SDS-PAGE and Western blot analysis. Immunoblotting was performed with anti-IFITM2 (not detected), anti-IFITM3, anti-IFITM1, and anti-actin. (C) Primary HSAEC were transfected with siRNA targeting IFITM3 or control siRNA for 48 hours. VSV-CoV-2 (50 μL) was added to cells and infection was measured by GFP expression at 24 hours post-infection using flow cytometry. siRNA-transfected cells were subjected to SDS-PAGE and Western blot analysis. Immunoblotting was performed with anti-IFITM2 (not detected), anti-IFITM3, anti-IFITM1, and anti-actin. (D) Semi-transformed nasal epithelial cells (UNCNN2TS) were treated with 20 μM Rap, Eve, Tem, Rid, or an equivalent volume of DMSO for 4 hours and whole cell lysates were subjected to SDS-PAGE and Western blot analysis. Immunoblotting was performed with anti-IFITM2 (not detected), anti-IFITM3, anti-IFITM1, and anti-actin. Immunoblots are representative of 3 independent experiments. Means and standard error were calculated from 4 experiments. Statistical analysis was performed with student's T test and asterisks indicate significant difference from control siRNA. *, $p < 0.05$; **, $p < 0.01$. Rel.; relative.

Supplemental Figure 2

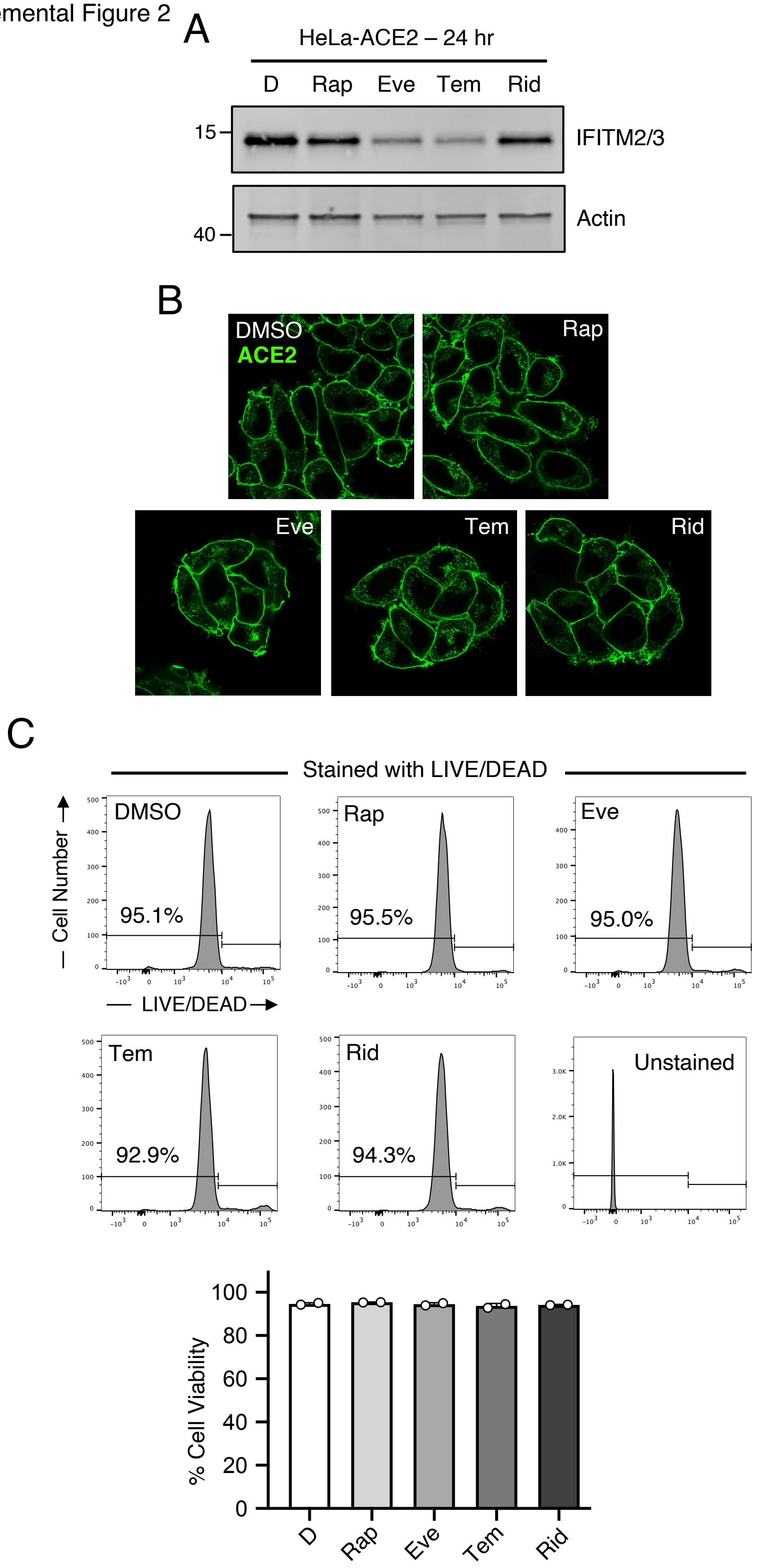

**Supplemental Figure 2:** (A) HeLa-ACE2 were treated with 20 μM Rap, Eve, Tem, Rid, or an equivalent volume of DMSO for 24 hours and whole cell lysates were subjected to SDS-PAGE and Western blot analysis. Immunoblotting was performed with anti-IFITM2/3 and anti-actin. (B) HeLa-ACE2 cells were treated with 20 μM Rap, Eve, Tem, Rid, or the equivalent volume of DMSO for 4 hours and cells were fixed, permeabilized, stained with anti-ACE2, and imaged by confocal immunofluorescence microscopy. Images represent a single, medial Z section. (C) HeLa-ACE2 cells were treated with 20 μM Rap, Eve, Tem, Rid, or the equivalent volume of DMSO for 4 hours and subsequently fixed and stained with LIVE/DEAD Fixable Red Dead Cell Stain Kit for 30 minutes according to manufacturer's instructions. Cells were analyzed by flow cytometry. Means and standard error were calculated from 2 experiments. All immunoblots are representative of three independent experiments.

Supplemental Figure 3

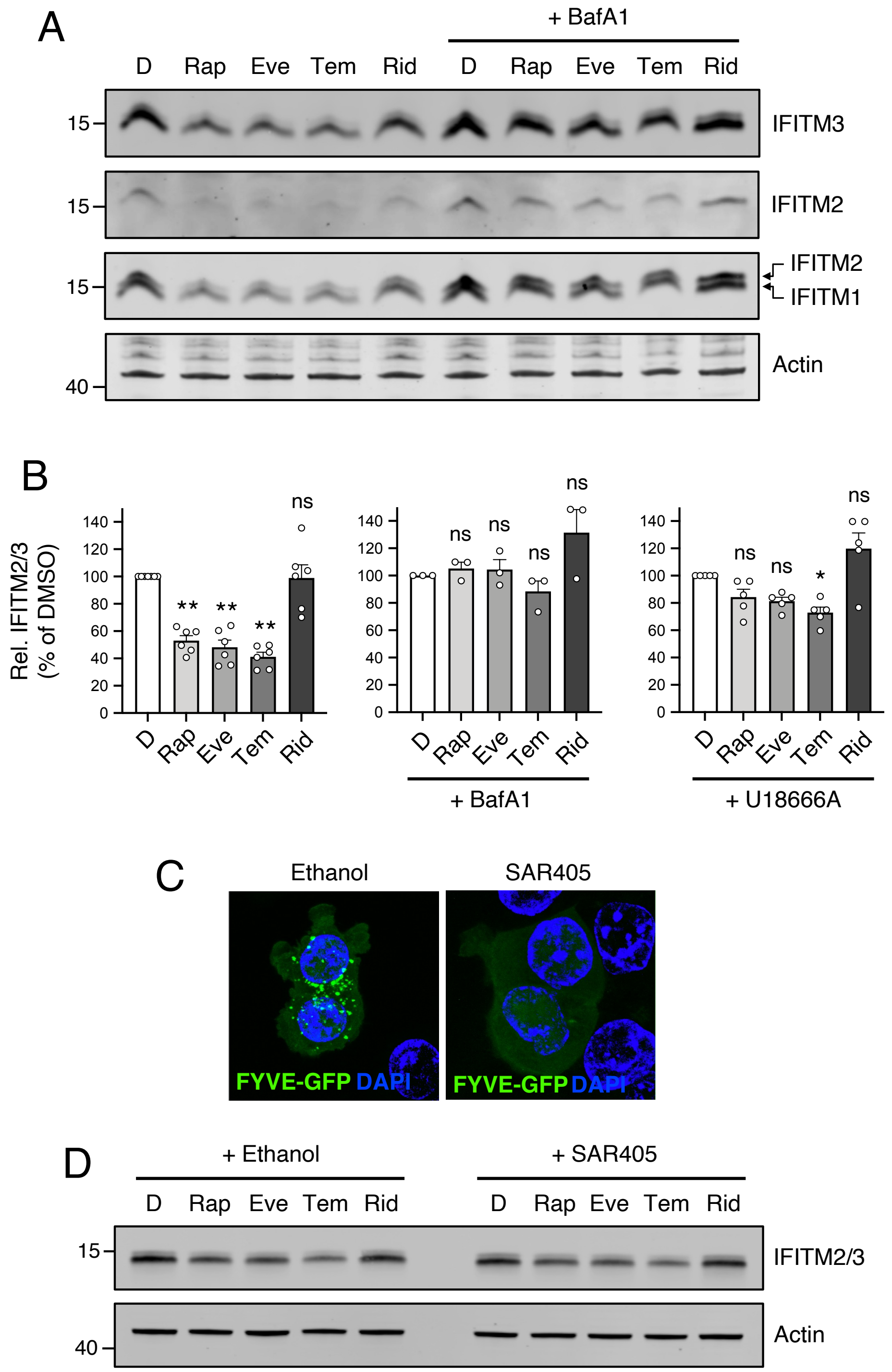

**Supplemental Figure 3:** (A) HeLa-ACE2 were treated with 20 μM Rap, Eve, Tem, Rid, or an equivalent volume of DMSO, in the presence or absence of 1 μM Bafilomycin A1, for 4 hours and whole cell lysates were subjected to SDS-PAGE and Western blot analysis. Immunoblotting was performed with anti-IFITM2, anti-IFITM1, anti-IFITM3, and anti-actin (in that order) on the same nitrocellulose membrane. (B) HeLa-ACE2 were treated with 20 μM Rap, Eve, Tem, Rid, or an equivalent volume of DMSO in the presence of 1 μM Bafilomycin A1, 5 μg/mL U18666A, or neither, for 4 hours. Cells were then fixed, permeabilized, and stained with anti-IFITM2/3. IFITM2/3 protein levels were measured using flow cytometry. Means and standard error were calculated from 3-6 experiments. (C) HeLa-ACE2 cells were transected with FYVE-GFP for 24 hours followed by treatment with 100 nM SAR405 or an equivalent volume of ethanol (vehicle) for 3 hours. Cells were fixed and imaged by confocal immunofluorescence microscopy. For each condition, a Z-stack of 25 slices is shown as a maximum intensity projection. (D) HeLa-ACE2 were treated with 20 μM Rap, Eve, Tem, Rid, or an equivalent volume of DMSO in the presence or absence of 100 nM SAR405 for 4 hours and whole cell lysates were subjected to SDS-PAGE and Western blot analysis. Immunoblotting was performed with anti-IFITM2/3 and anti-actin on the same nitrocellulose membrane. Statistical analysis was performed with one-way ANOVA and asterisks indicate significant difference from DMSO. *, $p < 0.05$; **, $p < 0.01$. ns; not significant. Rel.; relative.

Supplemental Figure 4

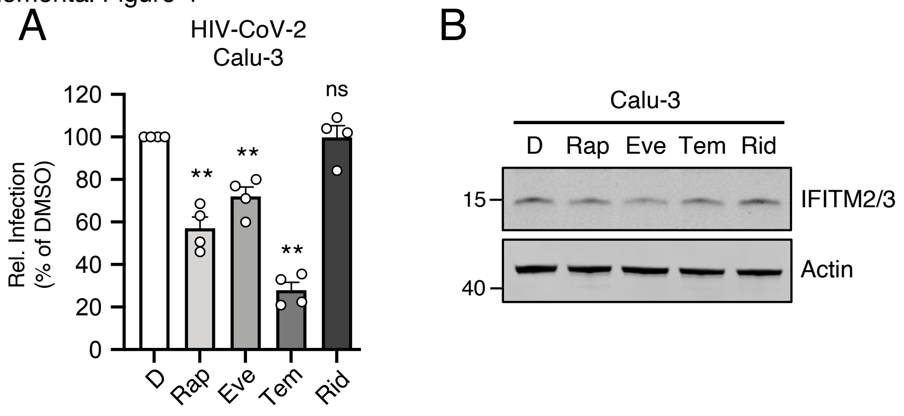


**Supplemental Figure 4:** (A) Calu-3 cells were treated with 20 µM Rap, Eve, Tem, Rid, or the equivalent volume of DMSO for 4 hours. HIV-CoV-2 (100 ng p24 equivalent) was added to cells and infection was measured by luciferase activity at 48 hours post-infection. Luciferase units were normalized to 100 in the DMSO condition. Means and standard error were calculated from 4 experiments. Statistical analysis was performed with one-way ANOVA and asterisks indicate significant difference from DMSO. *, $p < 0.05$; **, $p < 0.01$. ns; not significant. Rel.; relative. (B) Calu-3 cells were treated with 20 µM Rap, Eve, Tem, Rid, or the equivalent volume of DMSO for 4 hours. Whole cell lysates were subjected to SDS-PAGE and Western blot analysis. Immunoblotting was performed with anti-IFITM2/3 and anti-actin on the same nitrocellulose membrane.

Supplemental Figure 5

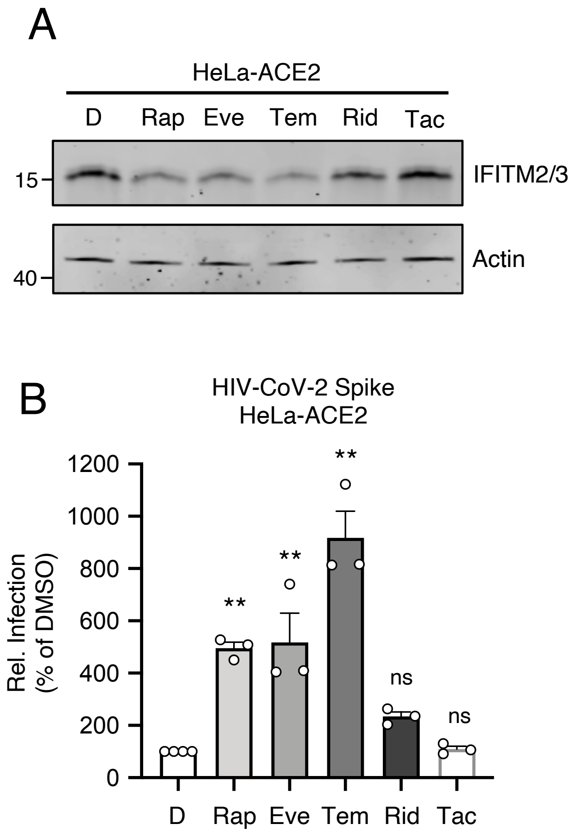


**Supplemental Figure 5:** (A) HeLa-ACE2 cells were treated with 20 µM Rap, Eve, Tem, Rid, Tac, or the equivalent volume of DMSO for 4 hours. Whole cell lysates were subjected to SDS-PAGE and Western blot analysis. Immunoblotting was performed with anti-IFITM2/3 and anti-actin on the same nitrocellulose membrane. (B) HeLa-ACE2 cells were treated with 20 µM Rap, Eve, Tem, Rid, Tac, or the equivalent volume of DMSO for 4 hours. HIV-CoV-2 (100 ng p24 equivalent) was added to cells and infection was measured by luciferase activity at 48 hours post-infection. Luciferase units were normalized to 100 in the DMSO condition. Means and standard error were calculated from 3 experiments. Statistical analysis was performed with one-way ANOVA and asterisks indicate significant difference from DMSO. *, $p < 0.05$; **, $p < 0.01$. ns; not significant. Rel.; relative.

Supplemental Figure 6

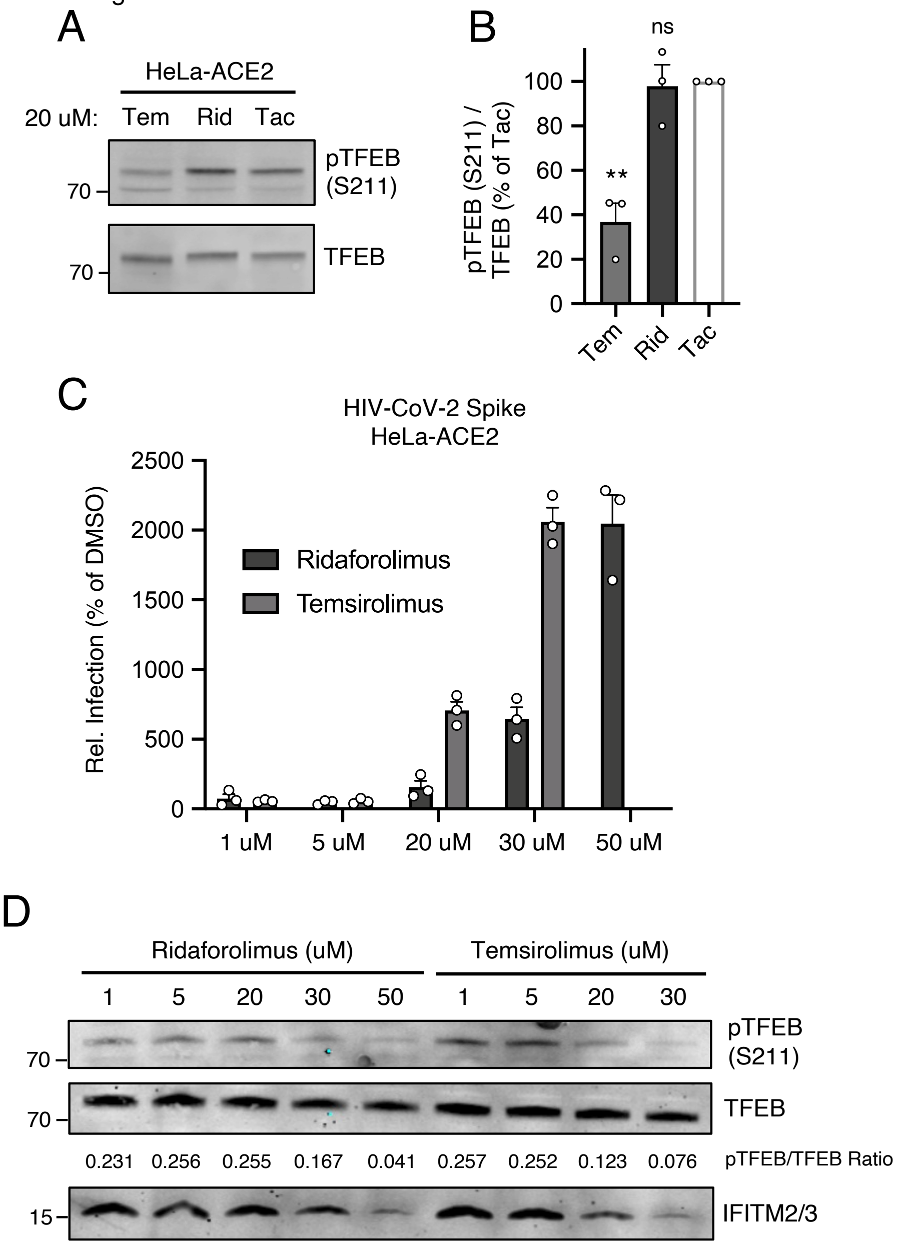


**Supplemental Figure 6:** (A) HeLa-ACE2 cells were treated with 20 µM Tem, Rid, or Tac for 4 hours. Whole cell lysates were subjected to SDS-PAGE and Western blot analysis. Immunoblotting was performed with anti-pTFEB (S211) and anti-TFEB on the same nitrocellulose membrane. (B) pTFEB (S211) levels were divided by total TFEB levels and summarized as an average of 3 experiments. (C) HeLa-ACE2 cells were treated with the indicated concentrations of Tem or Rid for 4 hours. HIV-CoV-2 (100 ng p24 equivalent) was added to cells and infection was measured by luciferase activity at 48 hours post-infection. Luciferase units were normalized to 100 in the DMSO condition. Means and standard error were calculated from 3 experiments. Rel.; relative. (D) HeLa-ACE2 cells were treated with the indicated concentrations of Tem or Rid for 4 hours and subjected to SDS-PAGE and Western blot analysis. Immunoblotting was performed with anti-pTFEB (S211), anti-TFEB, and anti-IFITM2/3 on the same nitrocellulose membrane. Levels of pTFEB (S211) were divided by total TFEB to obtain pTFEB/TFEB ratios.

Supplemental Figure 7

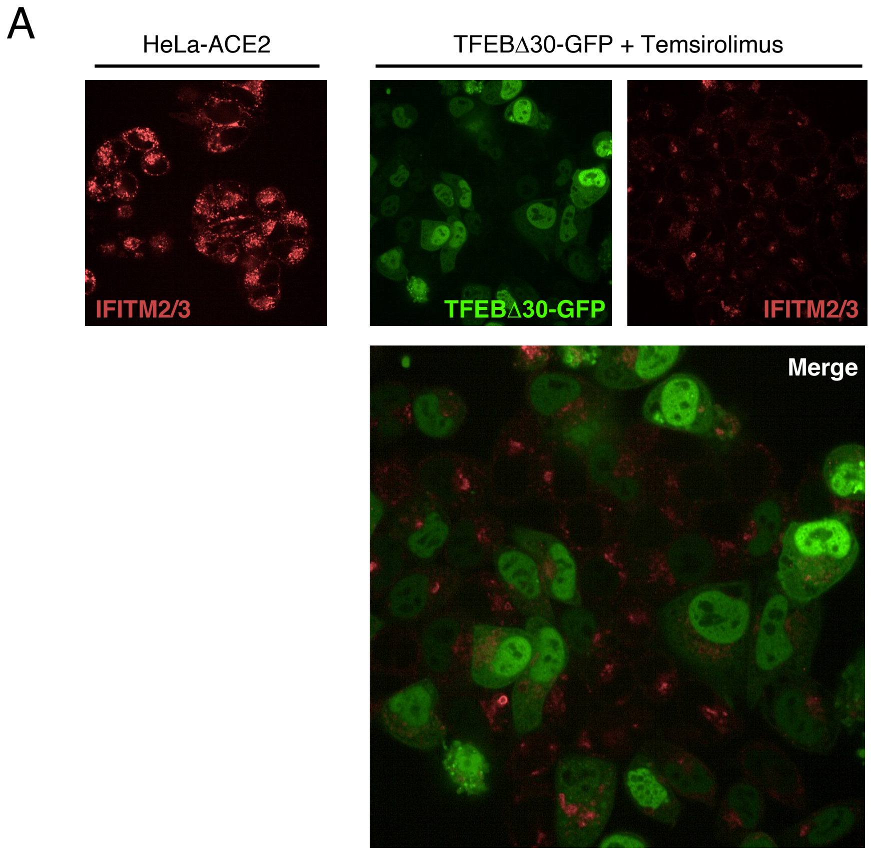


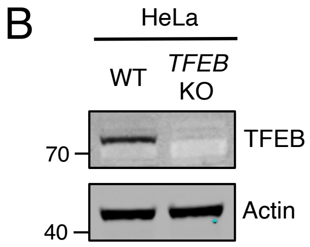


**Supplemental Figure 7:** (A) HeLa-ACE2 were transfected with 0.5 μg TFEBΔ30-GFP for 24 hours and treated with 20 μM Tem for four hours. Cells were then fixed, permeabilized, stained with anti-IFITM2/3, and imaged by confocal immunofluorescence microscopy. Representative images are shown and anti-IFITM2/3 staining in untreated HeLa-ACE2 are shown for comparison. (B) Whole cell lysates from HeLa WT and HeLa *TFEB* KO cells were subjected to SDS-PAGE and Western blot analysis. Immunoblotting was performed with anti-TFEB and anti-actin on the same nitrocellulose membrane.

Supplemental Figure 8

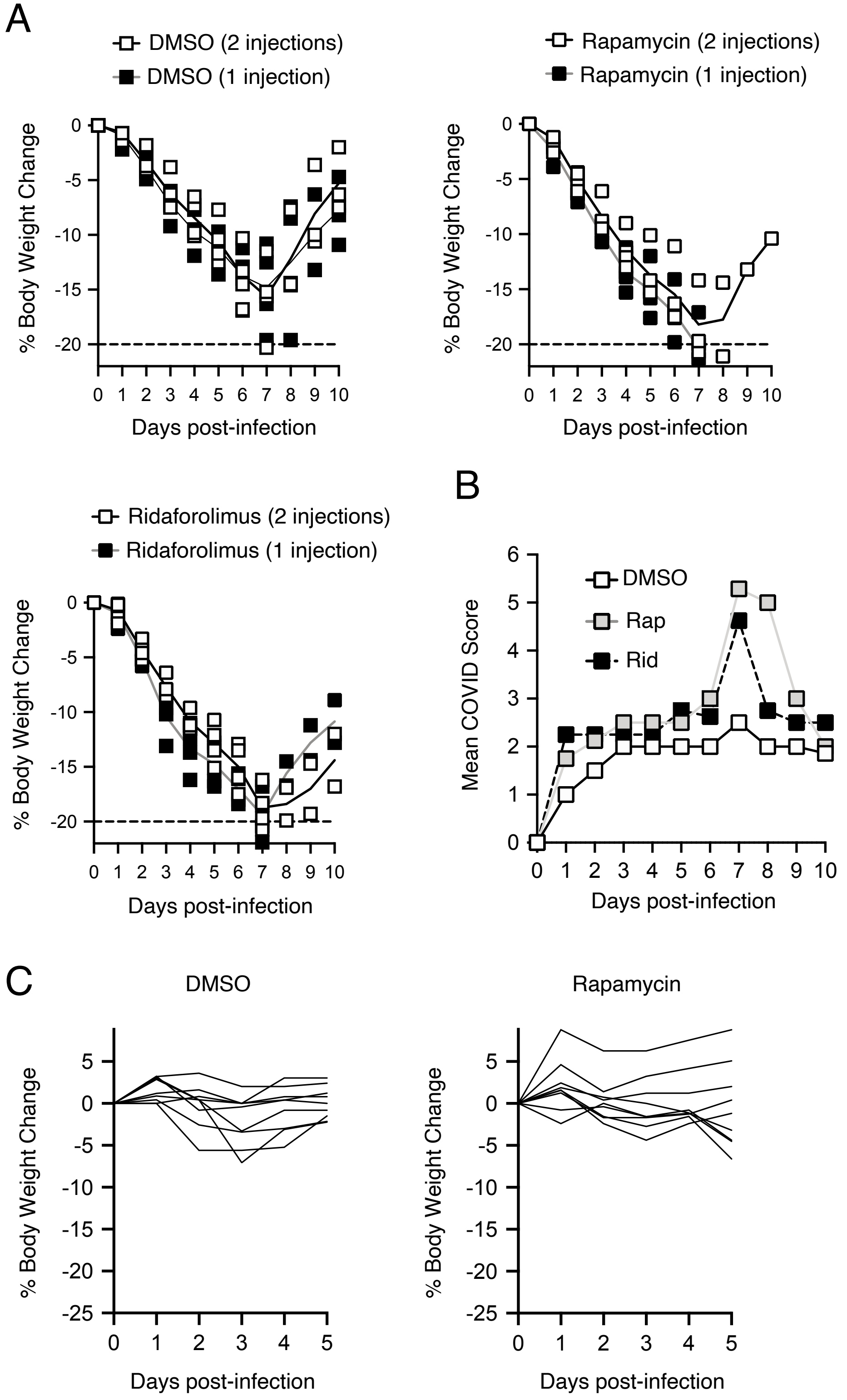

**Supplemental Figure 8:** (A) Body weight measurements for individual hamsters following injections with DMSO, Rap, or Rid are plotted by day post-infection and presented as % body weight change relative to Day 0. Hamsters receiving one injection of 3 mg/kg DMSO, Rap, or Rid prior to infection (n=4, 1 injection) are indicated by black squares, while hamsters receiving one injection prior to infection as well as a second injection of 3 mg/kg DMSO, Rap, or Rid at Day 2 post-infection (n=4, 2 injections) are indicated by white squares. The average daily weight change for each group is indicated by grey and black lines, respectively. If and when a hamster lost 20% or more of its body weight or exhibited agonal breathing, it was euthanized and body weight measurements were stopped. (B) Mean COVID scores were calculated for infected hamsters treated with DMSO, Rap, or Rid. Each hamster was observed daily and scores were calculated additively as follows: mild ruffling of fur (1 point), hunched posture (1 point), moderate/severe ruffling of fur (2 points), lethargic (3 points), more than 20% weight loss (4 points). (C) Body weight trajectories for individual C57BL/6 mice treated with DMSO (n = 9) or Rap (n = 9) and infected with MA-SARS-CoV-2 are plotted by day post-infection (up to 5 days).

**Supplemental Table 1: Overview of hamster lung pathology**

| Group | ID | Number of injections | Day of euthanasia | Basis for euthanasia | Gross lung observations | COVID score on day of euthanasia | Lung $TCID_{50}$ (per gram) | Lung IL-6 (pg/mL) |
|---|---|---|---|---|---|---|---|---|
| DMSO | 2527 | 2 (pre-, post-infection) | D10 | End of Expt. | Abnormal | 2 | BD | 168.7 |
| | 2528 | | D7 | Morbid (WL) | Severely hemorrhagic | 6 | 1,879,699 | 69.3 |
| | 2529 | | D10 | End of Expt. | Abnormal | 2 | BD | 123.8 |
| | 2530 | | D10 | End of Expt. | Abnormal | 2 | BD | 45.7 |
| | 2539 | 1 (pre-infection) | D10 | End of Expt. | Abnormal | 2 | BD | 168.7 |
| | 2540 | | D10 | End of Expt. | Abnormal | 1 | BD | 136.6 |
| | 2541 | | D10 | End of Expt. | Abnormal | 2 | BD | 162.4 |
| | 2542 | | D10 | End of Expt. | Abnormal | 2 | BD | 113.7 |
| Rap. | 2531 | 2 (pre-, post-infection) | D6 | Morbid (AB) | Moderately hemorrhagic | 7 | 3,340,565 | 51.5 |
| | 2532 | | D7 | Morbid (WL) | Severely hemorrhagic | 7 | 1,479,290 | 49.6 |
| | 2533 | | D8 | Morbid (WL) | Severely hemorrhagic | 7 | 49,104 | 56.9 |
| | 2534 | | D10 | End of Expt. | Abnormal | 2 | BD | 56.9 |
| | 2543 | 1 (pre-infection) | D7 | Found Dead | N/A | 6 | N/A | N/A |
| | 2544 | | D7 | Morbid (WL) | Severely hemorrhagic | 6 | 46,232 | 66.8 |
| | 2545 | | D7 | Morbid (WL) | Severely hemorrhagic | 6 | 585,607 | 36.3 |
| | 2546 | | D8 | Morbid (WL) | Severely hemorrhagic | 6 | 1,196,837 | 53.4 |
| Rid. | 2535 | 2 (pre-, post-infection) | D10 | End of Expt. | Abnormal | 3 | BD | 107.8 |
| | 2536 | | D8 | Morbid (AB) | Moderately hemorrhagic | 6 | 201,613 | 36.1 |
| | 2537 | | D8 | Morbid (WL) | Severely hemorrhagic | 6 | 52,705 | 67.0 |
| | 2538 | | D10 | End of Expt. | Abnormal | 2 | BD | 97.6 |
| | 2547 | 1 (pre-infection) | D7 | Morbid (WL) | Severely hemorrhagic | 7 | 47,722 | 69.5 |
| | 2548 | | D7 | Morbid (WL) | Severely hemorrhagic | 7 | 29,899 | 63.0 |
| | 2549 | | D10 | End of Expt. | Abnormal | 2 | BD | 172.8 |
| | 2550 | | D10 | End of Expt. | Abnormal | 3 | BD | 202.2 |

WL; weight loss. AB; agonal breathing. BD; below detection (1,667)

### APPENDIX

Microscopic lung pathology summarized by group (increased grade indicates increased severity)

DMSO:
Grade 1 edema, grade 3 hyperplasia, grade 1+ mixed or mononuclear inflammation. No difference between 1 vs 2 injections. Relative to other groups: Edema was decreased.

Rapamycin:
Grade 2 edema, grade 2 hyperplasia, grade 3 mixed or mononuclear inflammation. No difference between 1 vs 2 injections. Relative to other groups: hyperplasia was milder, inflammation was more severe.

Ridaforolimus:
Grade 2 edema, grade 3+ hyperplasia, grade 2+ mixed or mononuclear inflammation. No difference between 1 vs 2 injections.

Individual pathology reports

| Animal ID | Sex | Group |
|---|---|---|
| 2527 | M | DMSO - 2 injections |

Death: Scheduled, Terminal Sacrifice
Day of Death: D10
Date/Time: 06 Aug 2021 8:22 AM
Specimen: Left lung
Microscopic analysis:
- Hyperplasia - bronchiolo-alveolar, grade 3
- Inflammation, mixed cell - alveolar, interstitial, grade 1
- Inflammation, mononuclear cell - vascular/perivascular, grade 2
- Syncytial cell - grade 1

| Animal ID | Sex | Group |
|---|---|---|
| 2528 | M | DMSO - 2 injections |

Death: Unscheduled, Moribund Sacrifice
Day of Death: D7
Date/Time: 03 Aug 2021 8:20 AM
Specimen: Left lung
Microscopic analysis:
- Edema - perivascular, grade 1
- Hemorrhage - alveolar, grade 2
- Hyperplasia - bronchiolo-alveolar, grade 3
- Hypertrophy - mesothelium, grade 1
- Inflammation, mixed cell - alveolar, interstitial, grade 2
- Inflammation, mononuclear cell - vascular/perivascular, grade 1
- Syncytial cell - grade 1

| Animal ID | Sex | Group |
|---|---|---|
| 2529 | M | DMSO - 2 injections |

Death: Scheduled, Terminal Sacrifice
Day of Death: D10
Date/Time: 06 Aug 2021 8:22 AM
Specimen: Left lung
Microscopic analysis:
- Fibrosis - pleura, grade 1
- Hyperplasia - bronchiolo-alveolar, grade 4
- Inflammation, mixed cell - alveolar, interstitial, grade 2
- Inflammation, mononuclear cell - vascular/perivascular, grade 2
- Syncytial cell - grade 1

| Animal ID | Sex | Group |
|---|---|---|
| 2530 | M | DMSO - 2 injections |

Death: Scheduled, Terminal Sacrifice
Day of Death: D10
Date/Time: 06 Aug 2021 8:22 AM
Specimen: Left lung
Microscopic analysis:
- Fibrosis - pleura, grade 2
- Hyperplasia - bronchiolo-alveolar, grade 3
- Inflammation, mixed cell - alveolar, interstitial, grade 2
- Syncytial cell - grade 1

| Animal ID | Sex | Group |
|---|---|---|
| 2531 | M | Rapamycin - 2 injections |

Death: Unscheduled, Moribund Sacrifice
Day of Death: D7
Date/Time: 03 Aug 2021 8:49 AM
Specimen: Left lung
Microscopic analysis:
- Hemorrhage - alveolar, grade 1
- Hyperplasia - bronchiolo-alveolar, grade 2
- Inflammation, mixed cell - alveolar, interstitial, grade 3
- Syncytial cell - grade 1

| Animal ID | Sex | Group |
|---|---|---|
| 2532 | M | Rapamycin - 2 injections |

Death: Unscheduled, Moribund Sacrifice
Day of Death: D7
Date/Time: 03 Aug 2021 8:21 AM
Specimen: Left lung
Microscopic analysis:
- Edema - perivascular, grade 1
- Hemorrhage - alveolar, grade 2
- Hyperplasia - bronchiolo-alveolar, grade 2
- Hypertrophy - mesothelium, grade 1
- Inflammation, mixed cell - bronchoalveolar, interstitial, grade 3

| Animal ID | Sex | Group |
|---|---|---|
| 2533 | M | Rapamycin - 2 injections |

Death: Unscheduled, Moribund Sacrifice
Day of Death: D8
Date/Time: 04 Aug 2021 8:21 AM
Specimen: Left lung
Microscopic analysis:
- Edema - perivascular, grade 1
- Hyperplasia - bronchiolo-alveolar, grade 2
- Hypertrophy - mesothelium, grade 1
- Inflammation, mononuclear cell - vascular/perivascular, grade 1
- Inflammation, mononuclear cell - alveolar, interstitial, grade 3

| Animal ID | Sex | Group |
|---|---|---|
| 2534 | M | Rapamycin - 2 injections |

Death: Scheduled, Terminal Sacrifice
Day of Death: D10
Date/Time: 06 Aug 2021 8:22 AM
Specimen: Left lung
Microscopic analysis:
- Edema - perivascular, grade 1
- Fibrosis - pleura, grade 1
- Hyperplasia - bronchiolo-alveolar, grade 2
- Hypertrophy - mesothelium, grade 1
- Inflammation, mononuclear cell - vascular/perivascular, grade 1
- Inflammation, mononuclear cell - alveolar, interstitial, grade 3

| Animal ID | Sex | Group |
|---|---|---|
| 2535 | M | Ridaforolimus - 2 injections |

Death: Scheduled, Terminal Sacrifice
Day of Death: D10
Date/Time: 06 Aug 2021 8:22 AM
Specimen: Left lung
Microscopic analysis:
- Hyperplasia - bronchiolo-alveolar, grade 2
- Inflammation, mononuclear cell - alveolar, interstitial, grade 2

| Animal ID | Sex | Group |
|---|---|---|
| 2536 | M | Ridaforolimus - 2 injections |

Death: Unscheduled, Moribund Sacrifice
Day of Death: D8
Date/Time: 04 Aug 2021 8:21 AM
Specimen: Left lung
Microscopic analysis:
- Edema - perivascular, grade 1
- Fibrosis - mesothelium, grade 1
- Hemorrhage - alveolar, grade 2
- Hyperplasia - bronchiolo-alveolar, grade 3
- Inflammation, mixed cell - alveolar, interstitial, grade 3
- Inflammation, mononuclear cell - vascular/perivascular, grade 1
- Syncytial cell - grade 1

| Animal ID | Sex | Group |
|---|---|---|
| 2537 | M | Ridaforolimus - 2 injections |

Death: Unscheduled, Moribund Sacrifice
Day of Death: D7
Date/Time: 03 Aug 2021 8:21 AM
Specimen: Left lung
Microscopic analysis:
- Edema - perivascular, grade 1
- Hemorrhage - alveolar, grade 1
- Hyperplasia - bronchiolo-alveolar, grade 3
- Hypertrophy - mesothelium, grade 2
- Inflammation, mixed cell - bronchoalveolar, interstitial, grade 3
- Inflammation, mononuclear cell - vascular/perivascular, grade 1
- Syncytial cell - grade 1

| Animal ID | Sex | Group |
|---|---|---|
| 2538 | M | Ridaforolimus - 2 injections |

Death: Scheduled, Terminal Sacrifice
Day of Death: D10
Date/Time: 06 Aug 2021 8:22 AM
Specimen: Left lung
Microscopic analysis:
- Hyperplasia - bronchiolo-alveolar, grade 4
- Inflammation, mononuclear cell - vascular/perivascular, grade 1
- Inflammation, mononuclear cell - alveolar, interstitial, grade 2
- Syncytial cell - grade 1

| Animal ID | Sex | Group |
|---|---|---|
| 2539 | M | DMSO - 1 injection |

Death: Scheduled, Terminal Sacrifice
Day of Death: D10
Date/Time: 06 Aug 2021 8:22 AM
Specimen: Left lung
Microscopic analysis:
- Fibrosis - pleura, grade 1
- Hemorrhage - alveolar, grade 1
- Hyperplasia - bronchiolo-alveolar, grade 3
- Hypertrophy - mesothelium, grade 1
- Inflammation, mixed cell - alveolar, interstitial, grade 1
- Inflammation, mononuclear cell - vascular/perivascular, grade 1

| Animal ID | Sex | Group |
|---|---|---|
| 2540 | M | DMSO - 1 injection |

Death: Scheduled, Terminal Sacrifice
Day of Death: D10
Date/Time: 06 Aug 2021 8:22 AM
Specimen: Left lung
Microscopic analysis:
- Hyperplasia - bronchiolo-alveolar, grade 2
- Inflammation, mononuclear cell - vascular/perivascular, grade 1
- Inflammation, mononuclear cell - alveolar, interstitial, grade 2

| Animal ID | Sex | Group |
|---|---|---|
| 2541 | M | DMSO - 1 injection |

Death: Scheduled, Terminal Sacrifice
Day of Death: D10
Date/Time: 06 Aug 2021 8:22 AM
Specimen: Left lung
Microscopic analysis:
- Fibrosis - pleura, grade 1
- Hemorrhage - alveolar, grade 1
- Hyperplasia - bronchiolo-alveolar, grade 3
- Hypertrophy - mesothelium, grade 1
- Inflammation, mixed cell - alveolar, interstitial, grade 2
- Inflammation, mononuclear cell - vascular/perivascular, grade 1

| Animal ID | Sex | Group |
|---|---|---|
| 2542 | M | DMSO - 1 injection |

Death: Scheduled, Terminal Sacrifice
Day of Death: D10
Date/Time: 06 Aug 2021 8:22 AM
Specimen: Left lung
Microscopic analysis:
- Fibrosis - pleura, grade 1
- Hyperplasia - bronchiolo-alveolar, grade 3
- Inflammation, mononuclear cell - alveolar, interstitial, grade 1

| Animal ID | Sex | Group |
|---|---|---|
| 2543 | M | Rapamycin - 1 injection |

Found dead on 04 Aug 2021 - no lung pathology performed

| Animal ID | Sex | Group |
|---|---|---|
| 2544 | M | Rapamycin - 1 injection |

Death: Unscheduled, Moribund Sacrifice
Day of Death: D7
Date/Time: 03 Aug 2021 8:22 AM
Specimen: Left lung
Microscopic analysis:
- Edema - perivascular, grade 1
- Hemorrhage - alveolar, grade 1
- Hyperplasia - bronchiolo-alveolar, grade 3
- Hypertrophy - mesothelium, grade 1
- Inflammation, mixed cell - bronchoalveolar, interstitial, grade 3
- Inflammation, mononuclear cell - vascular/perivascular, grade 1

| Animal ID | Sex | Group |
|---|---|---|
| 2545 | M | Rapamycin - 1 injection |

Death: Unscheduled, Moribund Sacrifice
Day of Death: D7
Date/Time: 03 Aug 2021 8:22 AM
Specimen: Left lung
Microscopic analysis:
- Edema - perivascular, grade 1
- Hemorrhage - alveolar, grade 1
- Hyperplasia - bronchiolo-alveolar, grade 2
- Inflammation, mixed cell - bronchoalveolar, interstitial, grade 3
- Inflammation, mononuclear cell - vascular/perivascular, grade 2
- Syncytial cell - grade 1

| Animal ID | Sex | Group |
|---|---|---|
| 2546 | M | Rapamycin - 1 injection |

Death: Unscheduled, Moribund Sacrifice
Day of Death: D7
Date/Time: 03 Aug 2021 8:22 AM
Specimen: Left lung
Microscopic analysis:
- Hemorrhage - alveolar, grade 2
- Hyperplasia - bronchiolo-alveolar, grade 2
- Hypertrophy - mesothelium, grade 1
- Inflammation, mixed cell - bronchoalveolar, interstitial, grade 3
- Inflammation, mononuclear cell - vascular/perivascular, grade 2
- Syncytial cell - grade 1

| Animal ID | Sex | Group |
|---|---|---|
| 2547 | M | Ridaforolimus - 1 injection |

Death: Unscheduled, Moribund Sacrifice
Day of Death: D7
Date/Time: 03 Aug 2021 8:22 AM
Specimen: Left lung
Microscopic analysis:
- Edema - perivascular, grade 1
- Hemorrhage - alveolar, grade 1
- Hyperplasia - bronchiolo-alveolar, grade 3
- Hypertrophy - mesothelium, grade 1
- Inflammation, mixed cell - alveolar, interstitial, grade 3
- Inflammation, mononuclear cell - vascular/perivascular, grade 2
- Syncytial cell - grade 1

| Animal ID | Sex | Group |
|---|---|---|
| 2548 | M | Ridaforolimus - 1 injection |

Death: Unscheduled, Moribund Sacrifice
Day of Death: D7
Date/Time: 03 Aug 2021 8:22 AM
Specimen: Left lung
Microscopic analysis:
- Edema - perivascular, grade 1
- Hemorrhage - alveolar, grade 2
- Hyperplasia - bronchiolo-alveolar, grade 3
- Hypertrophy - mesothelium, grade 1
- Inflammation, mixed cell - alveolar, interstitial, grade 3
- Inflammation, mononuclear cell - vascular/perivascular, grade 1
- Syncytial cell - grade 1

| Animal ID | Sex | Group |
|---|---|---|
| 2549 | M | Ridaforolimus - 1 injection |

Death: Scheduled, Terminal Sacrifice
Day of Death: D10
Date/Time: 06 Aug 2021 8:22 AM
Specimen: Left lung
Microscopic analysis:
- Fibrosis - pleura, grade 2
- Hyperplasia - bronchiolo-alveolar, grade 3
- Inflammation, mononuclear cell - alveolar, interstitial, grade 2
- Syncytial cell - grade 1

| Animal ID | Sex | Group |
|---|---|---|
| 2550 | M | Ridaforolimus - 1 injection |

Death: Scheduled, Terminal Sacrifice
Day of Death: D10
Date/Time: 06 Aug 2021 8:22 AM
Specimen: Left lung
Microscopic analysis:
- Hyperplasia - bronchiolo-alveolar, grade 3
- Inflammation, mononuclear cell - vascular/perivascular, grade 2
- Inflammation, mononuclear cell - alveolar, interstitial, grade 2
